# Governed by surface amino acid composition: HIV capsid passage through the NPC barrier

**DOI:** 10.1101/2025.03.13.643050

**Authors:** Liran Fu, Shiya Cheng, Dietmar Riedel, Leonie Kopecny, Melina Schuh, Dirk Görlich

**Affiliations:** Department of Cellular Logistics, Max Planck Institute for Multidisciplinary Sciences, Göttingen, Germany; Department of Meiosis, Max Planck Institute for Multidisciplinary Sciences, Göttingen, Germany; Laboratory for Electron Microscopy, Max Planck Institute for Multidisciplinary Sciences, Göttingen, Germany; School of Basic Medical Sciences, Wuhan University, China

## Abstract

Nuclear transport receptors (NTRs) carry cargo across the permeability barrier of nuclear pore complexes (NPCs) - an FG phase condensed from disordered but cohesive FG repeats. This phase repels ‘normal’ macromolecules but allows NTR passage. When HIV infects non-dividing cells, its capsid is transported into nuclei not like cargo but crosses NPCs like NTRs. We now uncovered the molecular determinants of the capsid’s NTR behavior: The FG-binding pocket is insufficient. Hexameric and pentameric capsomers contribute. The highly exposed outer capsid surface is key. It lacks ‘FG-repulsive’ charged residues (K,D,E) that are very abundant on ‘normal’ protein surfaces. ‘FG-attractive’ residues dominate the capsid surface. Introducing FG-repulsive ones impedes FG phase-partitioning, NPC-targeting and NPC-passage of assembled capsids. Capsids are thus made FG phase-soluble by myriads of transient FG-attractive interactions originating from individual surface sidechains. We discuss that CPSF6 releases the capsid from NPCs by switching its surface from FG-attractive to FG-repulsive.

## Main

Nuclear pore complexes (NPCs) are embedded in the nuclear envelope (NE) to conduct nucleocytoplasmic transport^1-3^. This includes receptor-mediated import of nuclear proteins as well as export of mRNAs or newly assembled ribosomes^4-6^. NPCs are equipped with a barrier that can be rationalized as an FG phase^7-9^ – assembled from intrinsically disordered but cohesive FG repeat domains with numerous phenylalanine-glycine (FG) motifs. This barrier is selectively permeable to nuclear transport receptors (NTRs) along with captured transport substrates, while rejecting macromolecules that are not recognized as valid cargo.

Typical NPCs contain ∼12 different FG repeat domains anchored at distinct Nups. Of these, the FG domain of Nup98^10,11^ appears to be the most critical one for the NPC barrier^12^. It occurs in high copy number^13^. It is highly depleted of charged residues, poorly soluble in water and therefore readily phase-separating from aqueous solutions^8^. The resulting condensed FG phase shows essentially the same permeability properties as NPCs themselves^8,14^. It excludes inert macromolecules (such as mCherry), but dissolves NTRs to high partition coefficients. NTRs recognize and bind cargoes^4^. They promote barrier passage of a selected cargo by enhancing its solubility within the FG phase^7^. The phase behavior of ‘cohesive’ FG repeat domains and a phase-related transport selectivity have been documented in a range of experimental systems^8,12,15-20^. This includes the observation that NPCs become non-selectively permeable when the cohesive Nup98 FG domain is replaced by a non-selective one^12^.

The importin β superfamily represents the largest NTR class^4,21,22^, including nuclear import receptors (importins), exportins, as well as biportins. Asymmetric transport cycles driven by the RanGTPase system^23^ allow an active pumping of cargoes against a concentration gradient. Importins, for example, capture cargo in the cytoplasm, translocate it through NPCs, and release it into the nucleus upon RanGTP binding. While the permeability barrier retains imported cargo inside nuclei, importin·RanGTP complexes return to the cytoplasm, where GTP-hydrolysis releases Ran and allows the importins to import the next cargo molecule.

Importin β-type NTRs are 90-140 kDa α-solenoid proteins^22^. Other NTRs include NTF2 (the importer of Ran; ref.^24^), the Mex67·Mtr2 dimer (the main exporter of RNA; ref.^25,26^), and Hikeshi (the importer of Hsp70; ref.^27^). All NTRs bind FG repeats in a multivalent fashion^26,28-30^. This binding counteracts the hydrophobic cohesive interactions between repeats^31^ and thus allows an NTR to “melt” through the FG phase^32^. It was recently discovered that the HIV-1 capsid also functions as an NTR^33,34^.

HIV is the retrovirus that causes AIDS by infecting and eventually eliminating immune cells^35-37^. Its capsid initially encloses two copies of the HIV genomic RNA^38^, along with reverse transcriptase and integrase^39,40^. It is enveloped by a viral membrane that fuses with the plasma membrane during entry into a target cell^41^. This membrane fusion releases the capsid into the cytoplasm, where an influx of dNTPs into the capsid initiates reverse transcription^42^. The resulting vDNA is eventually integrated into a chromosome^43^. Simple retroviruses require (the open) mitosis to access chromatin. HIV and other lentiviruses, however, efficiently infect also non-dividing cells with intact NEs^44^, suggesting that an incoming HIV genome can somehow cross NPCs.

It was long believed that the 60 nm x 120 nm sized capsid was too large for NPC passage, that capsid uncoating and release of the HIV genome occurred in the cytoplasm, and that the capsid-free pre-integration complex was the species that is imported into the cell nucleus (discussed in ref.^45^). This view has recently been revised when still intact capsids were detected inside nuclei during the course of infection^46-50^, when cryo-EM tomograms showed incoming capsids trapped at NPCs^51^ and when the diameter of the NPC scaffold was shown to be larger in intact cells (∼60 nm; ref.^52^) than in the specimens earlier analyzed from isolated nuclear envelopes^53^. Finally, it was discovered that mature capsids behave like NTRs, efficiently partitioning into an FG phase and targeting NPCs without any *trans*-acting factors^33,34^. This NTR-like behavior required capsid assembly. These findings also resolved the problem that the 60 nm width of the NPC scaffold is just sufficient to accommodate a “naked” but not an importin-coated capsid.

CPSF6 is a host factor with a nuclear localization involved in capsid transport^54-56^. It contains a single, rather unusual FG ‘repeat’ that can dock into a pocket of the capsid^57,58^. It releases capsids from NPCs into the nucleoplasm and guides them to prospective chromosomal integration sites^46,47^.

Capsid assembly^59^ begins with the translation of the gag polyprotein (reviewed in ref.^60^), which comprises the MA (matrix protein), CA (p24 capsid protein), NC (nucleocapsid) and p6 modules. MA gets myristylated, binds lipids, and targets gag to the plasma membrane^61,62^. CA assembles into the immature capsid^63^ in an inositol phosphate-assisted manner^64^, while NC recruits HIV genomic RNA to the nascent virions^65^. P6 initiates virus-budding from the plasma membrane^66^. The switch to mature capsids is triggered by the HIV protease^67^, which cleaves gag into its individual proteins^68^. This switch involves major rearrangements in the capsomer structures^69^. This elaborate assembly sequence probably prevents the capsid from getting (mis-) targeted to NPCs already in virus-producing cells.

## Results

### Not only CA hexamers but also pentamers confer an efficient FG barrier entry

This study focuses on molecular features that allow mature HIV-1 capsids to pass the FG barrier of NPCs and thus deliver the viral genome into the nucleus. This passage should rely on interactions with FG repeats; and indeed a deep binding pocket between the subunits of the CA hexamers has been described which can accommodate the unique FG repeat of CPSF6 or a Nup153 FG repeat^56-58^. There, the sidechain amide of asparagine N57 of the CA protein engages in a double H-bond with the backbone of the FG peptide. The N57A mutation abrogates this stable binding^56,70,71^ and impedes NPC passage^33^ as well as the partitioning into a Nup98 FG phase (refs.^33,34^ and figures below). The capsid comprises not only hexamers but also pentameric capsomers^72^. Pentamers are thought not to bind FG repeats; a loss of CPSF6 FG-binding was documented and explained by an N57 pocket re-arrangement^73,74^.

Still, we wondered whether pentamers contribute to FG phase entry. To investigate this, we used the CA G60A, G61P double mutant to reconstitute 20 nm T1 capsid spheres, built solely of 12 pentamers^73^. For comparison, we also assembled capsids from CA wild-type^73,75^, yielding large (∼80 x160 nm) cone-shaped CLPs (capsid-like particles) dominated by hexamers, as well as 40 nm capsid spheres with 30 hexamers and 12 pentamers^76,77^. These capsid species were all labelled by 15% GFP fusion to the CA C-terminus that points to the capsid’s interior. Negative-stain electron microscopy (EM) confirmed proper assembly (Fig.1a).

As a proxy for the NPC barrier, we assembled an FG phase from Nup98-type, perfect GLFG 12mer repeats^78^. This phase accumulated CLPs and 40 nm capsid spheres to partition coefficients of 1000-2000 (Fig.1c-d) - consistent with our previous observations^33^. Surprisingly, the pentamer-only spheres behaved the same, reaching an intra-phase partition coefficient of ∼750 (Fig.1c-d). In FG phases with other common FG motifs (SLFG or FSFG), they partitioned to similarly high coefficients (Extended Data Fig.1). This partitioning was specific: unassembled protomers accumulated 300-600 times less; mCherry (our internal, inert control) or non-fused GFP remained completely excluded (with partition coefficients of ≤ 0.1).

### NPC-targeting of capsids can occur independently of the Nup358 CypH domain

In complementary assays with digitonin-semipermeabilized HeLa cells, we observed that not only CLPs and 40 nm spheres but also pentamer-only spheres target NPCs with very high efficiencies and then colocalize with an anti-Nup133 nanobody (Fig.1e; ref. ^79^).

NPC-targeting could be mediated by direct FG interactions and/or by the CypH (cyclophilin homology) domain of Nup358/ RanBP2, which is known to bind the ‘cyclophilin-binding loop” (loop 2) of the capsid^80^. To disentangle these potential contributions, we repeated the NPC-targeting experiment using the XTC-2 cell line^81^ from *Xenopus laevis* – a species, which (like all modern amphibians) has lost the CypH domain from its Nup358 (Extended Data Fig.2). Once again, we observed efficient capsid targeting to NPCs (Fig.1f). This indicates a highly efficient capsid entry into the FG phase of NPCs. This was also true for the pentamer-only capsids, confirming that CA pentamers mediate productive FG interactions. The experiment also showed that the capsid-FG phase interactions are independent of species-specific FG domain features.

### Crucial FG interactions by the N57-pocket of CA pentamers

The re-arrangement of the N57 FG-binding pocket in CA pentamers^73,74^ might imply that asparagine N57 may be irrelevant for the FG phase interactions of pentamer-only spheres. However, Fig.1g-h documents the opposite: The N57A mutation impeded FG phase entry not only of the hexamer-dominated CLPs and 40 nm spheres, but also of the pentamer-only spheres

- to the extent that only faint staining on the FG phase surface remained. Therefore, pentamers’ N57 pockets not only bind FG peptides, but their N57 sidechain amide also engages in energetically relevant H-bonds. The pentamer-only capsid thus behaves, by all criteria, like a transport receptor with a general FG-binding capability. The reported binding defect of pentamers^73,74^ appears to apply only to the sterically more constrained CPSF6 FG peptide (see also below, Fig. 7).

### FG phase surface-arrest phenotype

To immerse into the FG phase, the capsid must locally resolve cohesive interactions between FG repeats. This ‘melting’ is associated with a ΔG penalty^31^. Full entry into the FG phase therefore requires FG-capsid interactions to release more free energy than this penalty. If less free energy is released, capsids will remain at the phase surface, where ‘free’ FG motifs are directly exposed to the aqueous phase.

N57A mutant capsids show this ‘surface arrest’ phenotype with all FG phases tested (Fig.1g-h, Extended Data Fig.3e, and Supplementary Figure S1), indicating a thermodynamically relevant FG interaction defect and illustrating the relevance of the two N57-mediated H-bonds for locking the peptide backbone of FG peptides. These H-bonds are evident in experimental structures with CPSF6 and Nup153 FG peptides^57^, as well as in AlphaFold models of CA capsomers with captured GLFG, SLFG, or FSFG peptides (Extended Data Fig.3 and Supplementary Figure S1).

### The highly exposed capsid surface has a very unusual amino acid composition

The residual surface-binding of the N57 mutant capsid indicates, however, that the mutant has not lost all FG interactions and that other capsid features are also relevant. To identify additional contributions, we considered a completely different mode, namely collective interactions between the surface of a client and the FG phase. The underlying concept^8,32,82^ considers that (1) the FG phase is a solvent for its clients, (2) any sufficiently exposed residue on the client surface comes into contact with FG repeats when dissolved in the phase (with

∼400 mg/ml FG mass^31^), and (3) these contacts can be energetically favorable, neutral or unfavorable compared to contacts with water^14,83^.

Based on this concept, we previously engineered GFP to pass NPCs either very slowly or very rapidly, with extreme variants spanning a 15 000-fold difference in rate, and with a nearly perfect correlation between rates and FG phase partitioning^14^. This engineering exercise also uncovered an amino acid scale for FG interactions (Fig.2a): Negative charges (Asp and Glu) as well as lysine are strongly FG-repulsive, whereas exposed hydrophobic residues, cysteine, methionine, histidine and also arginine attract GFP (or other clients) into the FG phase.

The FG-repulsive residues Asp, Glu and Lys are abundantly exposed on the surface of soluble globular proteins (Fig.2). They provide topological information during the folding process and keep these globular proteins water soluble. They frequently occur in solvent-exposed parts of α-helices, β-sheets, and loops. On the protruding part of the outer surface of the HIV-1 capsid, however, they are extremely depleted (Fig.2b-e). In fact, loop 1 (the β-hairpin), loop 2 (the CypA-binding loop) and loop 3 do not contain a single Asp, Glu, or Lys (Fig.2c,e). This compositional bias is so striking that we suspected a connection to the NTR-like behavior of the capsid.

### Key to FG phase entry: the lack of FG-repulsive residues on the capsid surface

To explore this possible connection experimentally, we analyzed 13 additional capsid mutants that had previously been reported^54,84-87^. In brief, we assembled CLPs from the respective mutant CA proteins, validated capsid assembly by size exclusion chromatography and negative-stain EM (Extended Data Fig.4), and probed the Nup98 GLFG phase-entry of these CLPs (which had been filled with sinGFP4a to act as tracer for intact capsids).

Most mutants entered the FG phase as efficiently as wild-type CLPs (Fig.3a and Extended Data Fig.5a). No decrease in partitioning was observed when poorly exposed residues (other than N57) were mutated or when an exposed residue was changed to an equally or more hydrophobic one, examples being the G89V and P90A loop2 mutations, which were previously designed to abrogate interactions with cyclophilin A/ the CypH domain^87^. The same was true for the E45A, G116A, L136M, or R143A mutations.

The H87Q and R132K mutations caused a ∼50% reduction in partitioning. The A92E and G94D mutations in the highly exposed loop 2, however, showed a partition defect similarly strong as the N57A exchange. They arrested the capsids at the FG surface and reduced the partitioning to ∼5% of the wild-type level (Fig.3a-c). This is consistent with the fact that the negatively charged aspartic and glutamic acid residues belong to the most FG-repulsive surface features, which in turn can be explained by the energetically rather unfavorable contacts between negative charges and phenylalanines.

To explore the outer surface of the capsid more systematically, we first identified additional positions that could be mutated without compromising capsid assembly. For this, we again used size exclusion chromatography and negative stain electron microscopy to validate faithful capsid assembly (Extended Data Fig.4). ‘Stable capsid mutants’ were subsequently tested in transport assays (Fig.3d, and see below). A drastic drop in capsid-partitioning into the FG phase was evident for six individual mutations to an FG-repulsive glutamate (Fig.3d and Extended Data Fig.5b). These included the isosteric Q9E exchange at the tip of the β-hairpin of loop 1, the V86E, A88E, I91E, and M96E mutations at loop 2, as well as the P123E exchange at loop 3. Moreover, the R97K mutation impeded the FG phase partitioning. It preserves a positive charge; however, the two amino acids behave differently towards the FG phase^14^. Lysine is FG-repulsive. Arginine is FG-attractive, the explanations being (1) that its planar guanidinium group engages more readily in a cation-π interaction with phenylalanines of the FG repeat domain, (2) that it is an excellent hydrogen bond donor for carbonyl oxygen acceptors in sidechains and the polypeptide backbone (discussed in ref.^31^), and (3) that arginine is more readily transferred from an aqueous to a hydrophobic environment^31,83^. The partitioning defect of the R97K capsid mutant reflects these differences remarkably well.

The previously mentioned R132K mutant showed a similar trend, but its defect was much weaker, probably because R132 is less exposed (Extended Data Fig.6). This appears to be a general pattern: R143 is located even deeper in the inter-capsomer cavity, which explains why the R143K mutation had no effect on capsid-partitioning. Likewise, mutating leucine L136 at the bottom of the inter-capsomer cavity to glutamate (L136E) had no effect on phase entry, while the similar mutation of the highly exposed alanine 92 (A92E) was very detrimental (compared in Extended Data Fig.6). This non-equivalence emphasizes that the solubility of the HIV capsid in the hydrophobic FG phase is ruled by the surface properties of its protruding and highly exposed parts.

For the FG partitioning mutants studied here, one could argue that the observed effects are not due to the newly introduced FG-repulsive residues, but rather to the loss of the original ones. To address this, we next analyzed different substitutions for the same residue (Fig.3e and Extended Data Fig.5c). The mutations of histidine 87 in loop 2 to glycine (H87G) reduced the FG partitioning of the capsid only marginally. The H87Q exchange had a stronger but still mild effect. The H87E mutation, however, was detrimental. It reduced the partitioning ∼15-fold and left transport intermediates arrested at the FG phase surface. Thus, the defect cannot be explained by the loss of the imidazole moiety alone. What mattered was indeed the exchange to the FG-repulsive glutamate with its negatively charged sidechain. The same pattern was observed for mutations of V86, A88, I91, A92, and M96: Exchanges to glycine had only mild effects, while mutations to glutamate reduced the capsid-partitioning >10-fold and led to surface-arrested intermediates.

### Additive effects of capsid-partitioning mutations

Up to this point, we have identified 12 individual capsid mutants with a clear defect in FG phase partitioning. Yet, all of them retained some FG interactions. A combination of partitioning mutations, however, further reduced the FG phase signal of the capsids to the point of complete loss of partitioning and surface binding. This was the case when three FG-repelling surface mutations (H87Q+A92E+G94D) were combined with the N57A pocket mutation (Fig.4a and Extended Data Fig.7a).

Likewise, there is also a clear synergy between surface mutations alone, as seen by the greatly reduced signal of the Q9E+A92E double mutant CLPs (Fig.4b and Extended Data Fig.7b). The V86E+A88E+I91E+A92E quadruple mutant capsid combines four FG-repelling loop 2 mutations. It has lost all FG phase-partitioning and surface-binding. This is remarkable, because not only does the quadruple mutation keep the N57 pocket intact, but loop 2 does not even contact the FG repeat portion that docks into the N57 pocket - neither in the reported CA hexamer structures with bound CPSF6 or Nup153 FG peptides (PDB 4WYM and 4U0C) nor in structural models with other FG peptides (Extended Data Fig.3 and Supplementary Figure S1). Thus, the four negative charges dominate in their repulsion from the condensed FG phase – not only with respect to the still intact N57 pocket, but also with respect to all the other FG-attractive residues that still cover the outer surface of the capsid (e.g. H87, G94, M96, R97, or P123).

### Capsid mutations that impede FG phase-partitioning also interfere with NPC-targeting

In a next step, we compared the targeting of wild-type and mutant CLPs to NPCs of digitonin-semipermeabilized HeLa cells (Fig.5). To label intact capsids, we again loaded them with sinGFP4a. We not only reproduced the finding that the N57A pocket mutation reduced the NPC-targeting^33^, we also observed that all of the FG-repelling surface mutations had an at least equally deleterious effect. In fact, most of these mutations (e.g. A88E, A92E, or M96E) impeded capsid-binding to NPCs even more. The effects of these mutations were, again, additive. The V86E+A88E+I91E+A92E and the N57A+H87Q+A92E+G94D quadruple mutations reduced capsid-binding to NPCs to background levels.

### Large CLPs completely pass mouse oocyte NPCs when CPSF6 is overexpressed

We previously microinjected 40 nm capsid spheres into the cytoplasm of mouse oocytes^33^. These spheres passed NPCs and accumulated at intranuclear structures that probably represent nuclear speckles that are targeted by HIV during genuine infections^47^. When we repeated the experiment with large CLPs (covalently labelled with GFP), we observed only prominent binding to the nuclear envelope, respectively NPCs, but no intranuclear signal (Fig.6a, ‘WT’). This can be explained by a very slow NPC passage, because the same capsids showed prominent speckle-binding when injected directly into the nucleus.

We reasoned that capsid release from NPCs into the nucleoplasm was rate-limiting and that this slow step could be accelerated by increasing the concentration of the host factor CPSF6^46,54-56,88^. To test this, we microinjected mRNA encoding either mScarlet or a CPSF6-mScarlet fusion. Four hours later, GFP-labelled CLPs were microinjected into the cytoplasm (along with a microinjection marker). Another 30 minutes later, the oocytes were imaged by confocal laser scanning microscopy. This revealed efficient translation of the mRNAs: non-fused mScarlet was evenly distributed throughout nucleus and cytoplasm, sparing only the nucleolus, while the mScarlet-CPSF6 fusion was efficiently imported into the nucleus, giving a clean nucleoplasmic signal with a notable speckle enrichment.

Strikingly, when the oocyte nuclei had been supplemented with CPSF6, cytoplasmically injected CLPs not only bound to the nuclear envelope, but also passed NPCs and accumulated in nuclear speckles (Fig.6b). A complementary experiment revealed that the CPSF5·CPSF6 complex is a potent antagonist of capsid-partitioning into the FG phase (Fig.7a). This antagonizing effect was highly specific and completely lost when the singular FG motif of CPSF6 was mutated to a GG motif. Thus, CPSF6 appears to promote the completion of NPC passage by ‘extracting’ the capsid from the FG phase.

### FG-repelling surface mutations block NPC passage of capsids

To now test capsid mutations for NPC passage defects, we supplemented oocytes with unlabelled CPSF6 before microinjecting GFP-tagged wild-type or mutant CLPs into the cytoplasm. Wild-type CLPs reached the nuclear speckles, whereas the N57A mutant did not (Fig.6c). The mutant effect was confirmed by two controls. First, co-injected wild-type CLPs, labelled with mScarlet-I3, reached the speckles of the same oocyte. Second, N57A mutant CLPs also accumulated at the speckles when microinjected directly into the nucleus (Fig.6a). Thus, the N57A mutation causes a genuine NPC passage effect, even though capsid-targeting to the nuclear envelope was still prominent.

The above characterized V86E+A88E+I91E+A92E surface mutant combines four exchanges of hydrophobic residues for an FG-repulsive glutamic acid. After nuclear microinjection, it effectively bound speckles (Fig.6a). Following cytoplasmic microinjection, however, it failed to reach intranuclear structures. It did not even show any detectable binding to the nuclear envelope, indicating a very tight block in NPC-targeting and NPC-passage. This was obviously a consequence of the failure of these mutant capsids to partition into the FG phase (Figs. 4-5). Thus, the presence of FG-attractive residues and the striking absence of FG-repulsive residues in the capsid’s β-hairpin and its two exposed loops are key determinants of both, entering the permeability barrier of NPCs and passing through the pores completely.

## Discussion

The HIV-1 capsid has evolved into a nuclear transport receptor (NTR) that readily partitions into the FG phase-based permeability barrier of NPCs. It serves as a cargo container for the encapsulated genetic material, shielding it from anti-viral sensors in the cytoplasm and delivering it through an intact nuclear envelope into the nucleus, thus allowing the infection of non-dividing cells.

### Both hexameric and pentameric capsomers contribute to FG phase-partitioning

The capsid is very large – at the very limit of what can possibly pass through an NPC scaffold - and is typically made up of 200-250 hexameric and 12 pentameric capsomers^72,89-93^. It was previously thought that only the hexamers engage in FG interactions^73,74^. However, we have now found that pentamers (represented by pentamer-only T1 spheres^73^) mediate a very efficient FG phase entry as well (Fig.1). Indeed, CA hexamers and pentamers appear very similar in their FG interactions: Efficient NPC-targeting and FG phase-partitioning of either species require assembly into a capsid structure (Fig.1c-f). Furthermore, point mutations that impede the FG-partitioning of hexamer-dominated capsids also compromise the partitioning of the pentamer-only T1 capsid. This applies to the N57A mutation as well as to the surface loop mutations (Fig.1g, Fig.4c, and Extended Data Fig.7c, Extended Data Fig.8a-b).

**Fig. 1.**
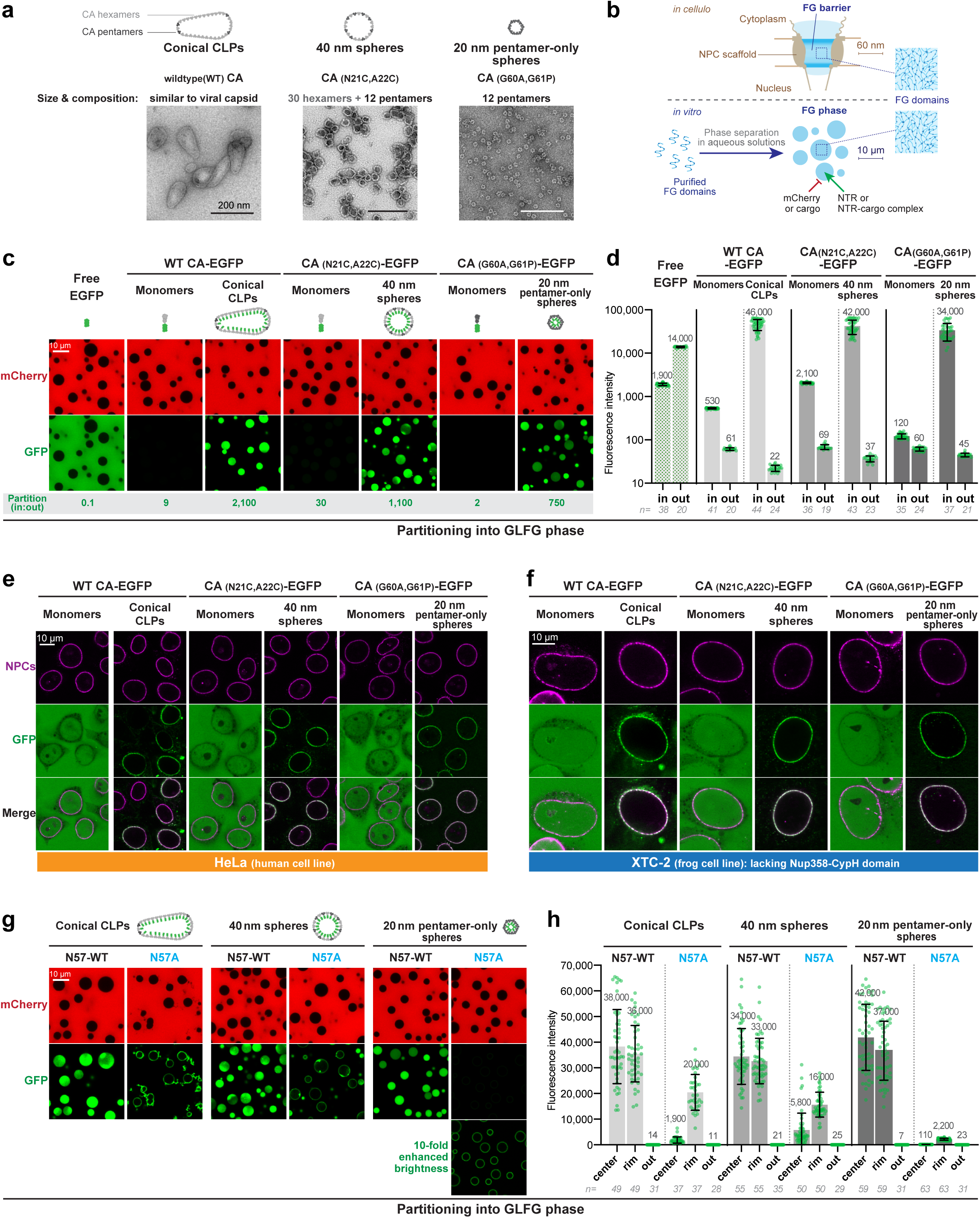
Hexameric and pentameric capsomers mediate FG phase-partitioning and NPC-targeting. **a,** Negative stain electron micrographs of the HIV-1 capsid species used. **b,** Concept of the FG phase as a permeability barrier of NPCs, reconstitution of such a barrier by phase separation of barrier-forming cohesive FG domains, and testing of its transport properties. **c,** A Nup98-type FG domain comprising 52 perfect 12mer GLFG repeat units (“prf.GLFG_52×12_”)^78^ was allowed to phase separate, forming FG phases of near-spherical shape. Indicated fluorescent species were subsequently added and detected one hour later by confocal laser scanning microscopy (CLSM). Free mCherry and EGFP remained well excluded from the FG phase, with partition coefficients (in: out) of ≤0.1. Large CLPs, 40 nm spheres, and 20 nm T1 ‘pentamer-only’ spheres were each labeled by ∼15% CA with a C-terminally fused EGFP, pointing to capsids’ interior. They accumulated essentially completely inside the FG phase, reaching partition coefficients of ∼1000. This was 30-300 times higher than the corresponding monomeric CA-EGFP fusions and 10 000 times higher than free EGFP. **d,** Quantification of **c** but with larger datasets. Datapoints are the integrated inside (in) signal of one FG particle or that of a corresponding outer (out) region. Numbers are means; bars indicate mean ± s.d.; n = number of quantified FG particles and outer areas, respectively. **e, f,** Specific targeting of capsid species to NPCs. Human HeLa cells or frog XTC-2 cells (whose Nup358 lacks a CypH domain) were grown on coverslips, treated with digitonin to permeabilize their plasma membranes, and incubated with the indicated fluorescent species. Confocal scans were taken directly through the live samples 30 min later. Laser settings were individually adjusted. NPCs were detected with an Alexa647-labelled anti-Nup133 nanobody^79^. Note that conical CLPs, 40 nm capsid spheres, and pentamer-only spheres showed highly efficient NPC-targeting in both cell types – comparable to nanobody-staining. By contrast, unassembled CA monomers showed no NPC enrichment and were evenly distributed throughout the cells. **g, h,** FG phase-partitioning of all three capsid species is strongly reduced by the N57A mutation. Experiments and quantifications were performed as in **c, d** but signals at the phase surface (rim) and the center were integrated separately (see also below, Fig.3b,c). Scan settings were identical for each wildtype-N57A mutant pair. Experiments were repeated independently with identical outcomes (*N* = 4). Scale bars, 200 nm in **a**, 10 μm in **c, e, f,** and **g**.

Although low in count, the contribution of pentamers is probably quite relevant, because they are concentrated at the capsid’s narrow end, which appears to insert first^51^ into NPCs and which should therefore have a particularly high FG-partitioning propensity.

The condensation of FG repeat domains into an FG phase is driven by cohesive interactions that are of a hydrophobic nature^15,31,32,82,94^. Partitioning of a client into this phase requires a local ‘melting’ of these cohesive interactions. This easily happens when the client (e.g. an NTR) binds to (the hydrophobic) FG motifs. Indeed, NTRs belong to the most hydrophobic soluble proteins^82^. This poses a fascinating design challenge, namely how to make a protein surface ‘NTR-like hydrophobic’ without causing aggregation and non-cognate interactions?

### FG-binding pockets and FG-wettable surface

Binding pockets for FG motifs are one solution. They have been identified by X-ray crystallography and cryo-EM in several cellular NTRs, such as importin β, Xpo1/ CRM1 or the Mex67-Mtr2 dimer^26,28,30,95,96^. FG-binding pockets can confer excellent binding specificity without fully exposing their hydrophobicity. They contain mostly hydrophobic, but also polar residues, whose aliphatic moieties then engage in hydrophobic interactions with FG motifs, while their polar head groups hydrogen-bond, e.g., to the FG repeat backbone^26,28,96^. FG-binding pockets appear to confer rather strong FG interactions that can overcome the FG-repelling effects of the typically strong negative charge of cellular NTRs (here, the negative charge is required to transport also highly positively charged cargoes such as histones). It seems reasonable to assume that these details of FG interactions have evolved to optimize cargo transport.

The HIV-1 capsid has the N57 pocket which occurs in very high multiplicity (given the capsid’s repetitive architecture). It is located between adjacent CA monomers and is much deeper than FG-binding pockets in cellular NTRs. It uses the sidechains of L56, M66, L69, I73, and the aliphatic portion of K70 to form hydrophobic contacts with the FG motif phenylalanine. The sidechain amide of asparagine N57 engages in a double or triple hydrogen-bond with the backbone of the inserted FG repeat (ref.^57^; Extended Data Fig.3 and Supplementary Figure S1). N57 and L56 are absolutely conserved amongst HIV-1, HIV-2 and SIV (Extended Data Fig.9). K70, M66, L69, and I73 are also conserved, but show some conservative exchanges (K➔R, M➔L, L➔I/V, I➔V; Extended Data Fig.9). This suggests an ancestral function of this pocket and possibly a common NPC passage mechanism of lentiviral capsids (see also below).

However, this pocket is only one part of the story. The entire accessible outer surface of the mature capsid appears to be optimized for “wetting” by the FG phase (Figures 1-4). It has a highly unusual amino acid composition that follows a previously identified scale for FG phase entry^14^. The key feature is the complete absence of the FG-repulsive residues Asp, Glu and Lys in the exposed loops. This is a novel principle of encoding topogenic information. It is relevant because the introduction of repulsive glutamates impedes FG phase-partitioning, NPC-targeting and also NPC passage (Figures 3-6) - up to a complete block when mutations are combined. We have documented these phenotypes at nine positions: Q9 (loop1), V86, A88, H87, I91, A92, G94, M96 (loop2) and P123 (loop 3). A similar phenotype was observed for the R97K exchange, where an FG-attractive arginine was replaced by an FG-repulsive lysine. None of these mutations have been observed in HIV-1 isolates from patients (Table 1), suggesting that they are detrimental to viral fitness. Nevertheless, we assume that they still allow infection of proliferating cells, in which a capsid passage through NPCs is not a strict requirement.

**Table 1.** Circulating HIV strains strictly avoid FG-repelling capsid mutations.

| **Mutation** | **Position** | **FG phase-partitioning** | **NPC-targeting** | **Residues in naturally occurring strains** |
| --- | --- | --- | --- | --- |
| **N57A** | FG pocket | Severe defect | Reduced | N |
| **V86E** | loop 2 | Severe defect | Reduced | V, A, P, I, Q, T, M, L |
| **H87E** | loop 2 | Severe defect | Reduced | H, Q, P |
| **A88E** | loop 2 | Severe defect | Reduced | A, V |
| **I91E** | loop 2 | Severe defect | Reduced | I, V, N, A, F, H, L, P, Y, T, Q, M |
| **A92E** | loop 2 | Severe defect | Reduced | A, P, Q, V |
| **G94D** | loop 2 | Severe defect | Reduced | G, N |
| **M96E** | loop 2 | Severe defect | Reduced | M, I, L, V |
| **R97K** | loop 2 | Reduced | Reduced | R |
| **Q9E** | loop 1<br>(β-hairpin) | Severe defect | Reduced | Q, H, A |
| **P123E** | loop 3 | Reduced | Reduced | P, A, S, V, G |

One would expect enhanced FG phase-partitioning if capsid surface residues were changed to more FG-attractive ones. Such an effect is not obvious with our standard GLFG phase that already fully absorbs the wild-type CLPs. However, such an enhancement is evident for a stricter FSFG phase, which does not allow full entry of the wild-type CLPs. Here, the G89V mutant capsid enters four times more effectively (Extended Data Fig.8c-d).

N57 engages in a polar (H-bond) interaction with the FG repeat backbone, while the capsid surface residues analyzed here probably contribute mostly hydrophobic contacts. The different qualities of these interactions become experimentally apparent when the salt concentration is varied: Wild-type CLPs enter the GLFG phase at low (50 mM), medium (150-250 mM) or high salt (600 mM NaCl) concentrations (Extended Data Fig.10). The N57A mutant fails to partition at low and medium salt, but enters the phase at 600 mM NaCl, probably because this strengthens the remaining hydrophobic capsid surface-FG interactions. X➔E surface mutations, however, could not be rescued by high salt. In fact, the Q9E mutant even shows an inverse salt effect.

### A hydrophobic and yet aggregation-resistant capsid surface

The three outer loops are dominated by FG-attractive residues. The most attractive ones, tryptophan, tyrosine, and phenylalanine, however, are suspiciously missing (Fig.2, Table 1). Why is this? Perhaps, because highly exposed tryptophans or tyrosines are very prone to promiscuous interactions. We observed this in our previous engineering study on transforming GFP into an NTR^14^: Placing additional tryptophans or tyrosines onto the GFP surface greatly enhanced FG phase-partitioning and the rate of NPC passage, but also conferred massive aggregation with nuclear and cytoplasmic structures.

**Fig. 2.**
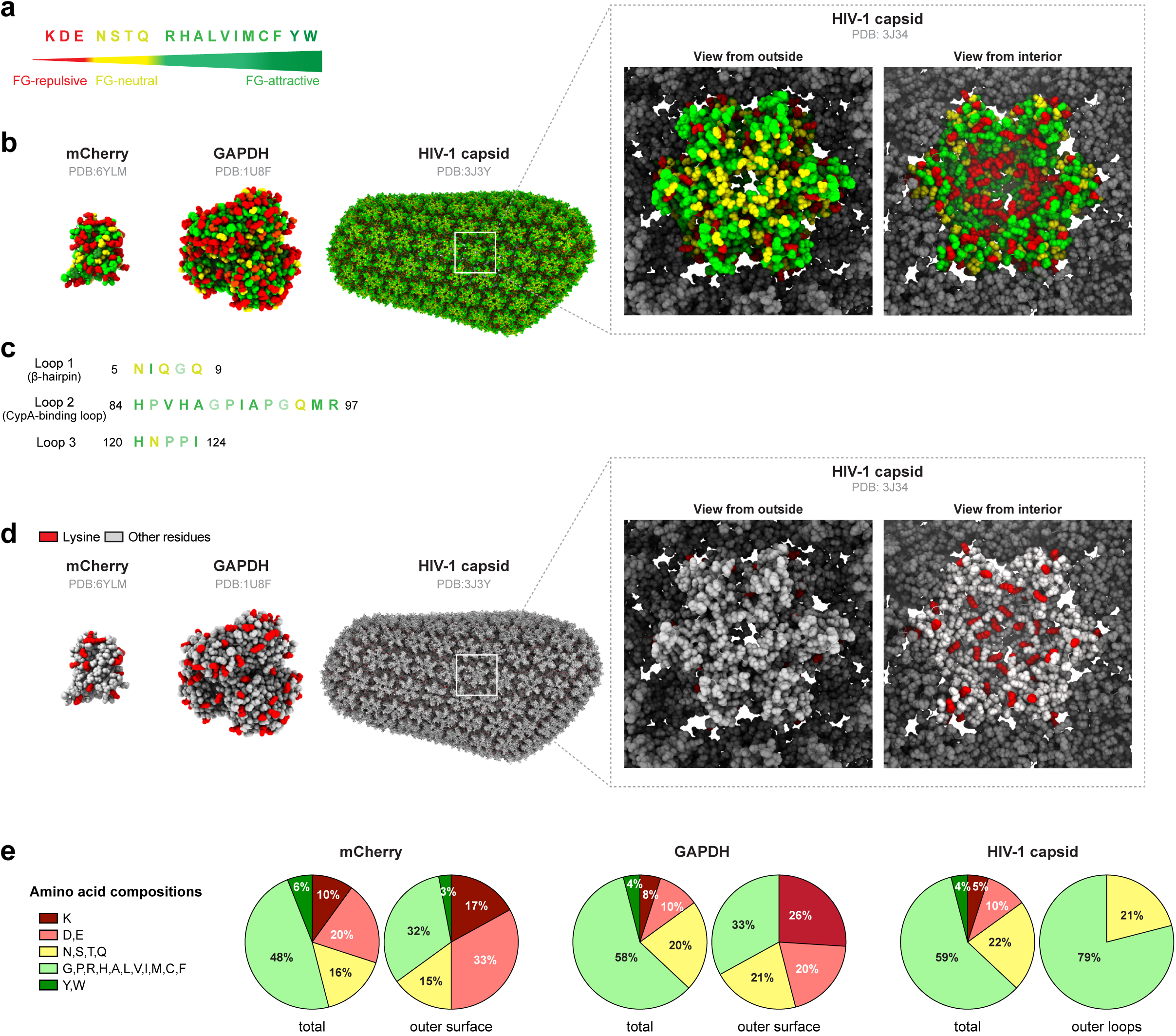
Surface amino acid composition of the HIV-1 capsid. **a,** A surface amino acid scale for partitioning into an FG phase^14^. FG-repulsive residues are coloured in red, FG-neutral residues in yellow, and FG-attractive residues in green or dark green. **b,** Comparison of the mCherry, GAPDH and HIV-1 capsid surfaces with residues coloured according to the FG phase-partitioning scale. Note that FG-repulsive residues (shown in red) are very abundant on the surface of mCherry, GAPDH (an example of cytoplasmic mass proteins), and on the inner surface of the capsid; but they are absent from protruding parts of the outer capsid surface. There are only a few in remaining outer surface, and these are not well accessible but mostly salt-bridged to FG-attractive R and H (E113-R97; E98-H84/ H87). **c,** Amino acids on the exposed loops of HIV-1 capsid are either FG-attractive or FG-neutral. **d,** Comparison of lysines exposed on the surfaces of mCherry, GAPDH and HIV-1 capsid. **e,** Diagrams compare total and surface amino acid compositions, classified by the scale of **a**.

By rational design, followed by mutagenesis and screening of many variants, we eventually obtained highly NPC-/ FG phase-specific and rapidly translocating NTR-like GFP variants^14^; but strikingly, they all lacked fully exposed tyrosines and tryptophans. It is remarkable that the evolution of the capsid into a specific NTR has arrived at a similar solution.

This must have occurred under strong selective pressure, since aggregation propensity scales with size, and thousands of cytoplasmic/ nuclear proteins were potential aggregation partners. There are fascinating parallels here to an antibody response by the immune system, which not only aims for strong target-binding but also selects *against* broad cross-reactivity with myriads of undesired (self-) targets. CDRs of early stage-antibodies often contain exposed tyrosines, perhaps because tyrosine-containing CDRs have such a high propensity to engage in interactions^97^. In well affinity-matured antibodies, however, tyrosines are typically replaced by more sophisticated and target-specific hydrophobic arrays^98^. In this sense, we consider the HIV-1 capsid to be a highly selectivity-matured entity – a masterpiece of evolution, created through very deep sampling of sequence space and the “testing” of extremely large numbers of variants during infection.

### Evolutionary pressure against surface lysines

In order to pass NPCs and thereby provide the selective advantage of infecting non-dividing cells, the capsid had to be highly optimized. Could this have evolved in a single step? Or did some other evolutionary pressure pave the way first? Possibly. At least as far as the suspicious absence of accessible lysines is concerned. Lysines are not only FG-repulsive, they are also acceptors for ubiquitin conjugation. We now propose that their elimination from the accessible outer surface (Fig.2d) came first – to protect incoming capsids from proteosomal degradation. Indeed, even the CA N-terminus is refractory to ubiquitin modification, since it is a secondary amine (proline) and is buried in the mature capsid^90,91,93^.

### FG-attractive capsid features are more ancient than HIV

The N57 FG-binding pocket is well conserved amongst lentiviruses, as are the outer capsid loops, which have a similar amino acid composition as HIV-1 (see above, Table 1, and Extended Data Fig.9). This suggests that lentiviral capsids share the ability to partition into the FG phase and pass NPCs, which would explain why lentiviruses are generally able to infect non-dividing cells. Indeed, our preliminary data indicate that also HIV-2 and SIV capsids can target the NPC autonomously and partition into an FG phase – though not quite as efficiently an HIV-1 M group capsid.

In contrast, capsids of simple retroviruses (such as the mouse leukemia virus MLV^99,100^) lack the FG-attractive extension of loop 2. This suggests that they cannot pass NPCs in the same way as HIV, which would explain why they infect quiescent cells only with marginal efficiency^44,101^.

### Escape from the energy well of the FG phase

HIV-1 capsids partition so strongly into the FG phase that the outside signal becomes undetectable (refs.^33,34^; Figs 1,3,4, and Extended Data Figures 1,3,5-8,10). Capsids are thus trapped in a deep energy well. How can they escape? During cellular transport, the RanGTPase system releases cargo from importin β-type NTRs and thus also from the NPC barrier (by nuclear RanGTP binding to importins or cytoplasmic GTP hydrolysis in cargo·exportin·RanGTP complexes^4,78^). The capsid, however, is not a RanGTP effector. HIV therefore needs a different strategy to release its genetic material from NPCs. Three come to mind:

1. Destruction of NPCs. Capsids can crack NPCs^102^. Yet, cracks are unlikely to trigger capsid escape from the FG phase, because the NPC scaffold elements and thus the FG mass stay in place. Furthermore, such cracks do not occur in all infectable cell types. We regard them as ‘collateral damage’ and not as a requirement for capsid passage through NPCs. Anyway, it appears that the nucleocytoplasmic barrier of mouse oocytes has not ‘suffered’ from cracks. It continued to exclude our injection marker even after large numbers of capsids had inserted into and passed through NPCs (Fig.6).
2. Disassembly of still NPC-trapped capsids, leading to vDNA release. Since FG phase-partitioning is favored by capsid assembly (refs.^33,34^; Fig.1 and Extended Data Fig.1), the inverse should also hold true, namely capsid stabilization by immersion into the FG phase. However, once reverse transcription has sufficiently increased the volume of the enclosed nucleic acids^103,104^, also FG phase-trapped capsids will burst and release the viral genome into either the nucleus or the cytoplasm. This pathway likely permits infection if capsids (or capsid mutants) fail to interact with CPSF6 (see ref.^86^ and below).
3. The most plausible mechanism for a release of intact capsids from NPCs is an energy input from a nuclear binding reaction that disengages the FG nucleoporin repeats from the capsid. Such a mechanism would allow for a directional capsid release into the nucleoplasm (Fig.7). Indeed, CPFS6 is the perfect candidate for terminating the capsid’s NPC passage. It is a host factor for infection; it binds the capsid directly, it participates in the nuclear events during the establishment of infection^54-56,88^; and capsids accumulate at NPCs when CPSF6 is depleted^46^. We have now directly demonstrated that the partitioning of fully assembled capsids into the FG phase is potently antagonized by the CPSF5·CPSF6 complex and that the CPSF6 paralog CPSF7 has a similar albeit weaker effect (Fig.7a,b).

**Fig. 3.**
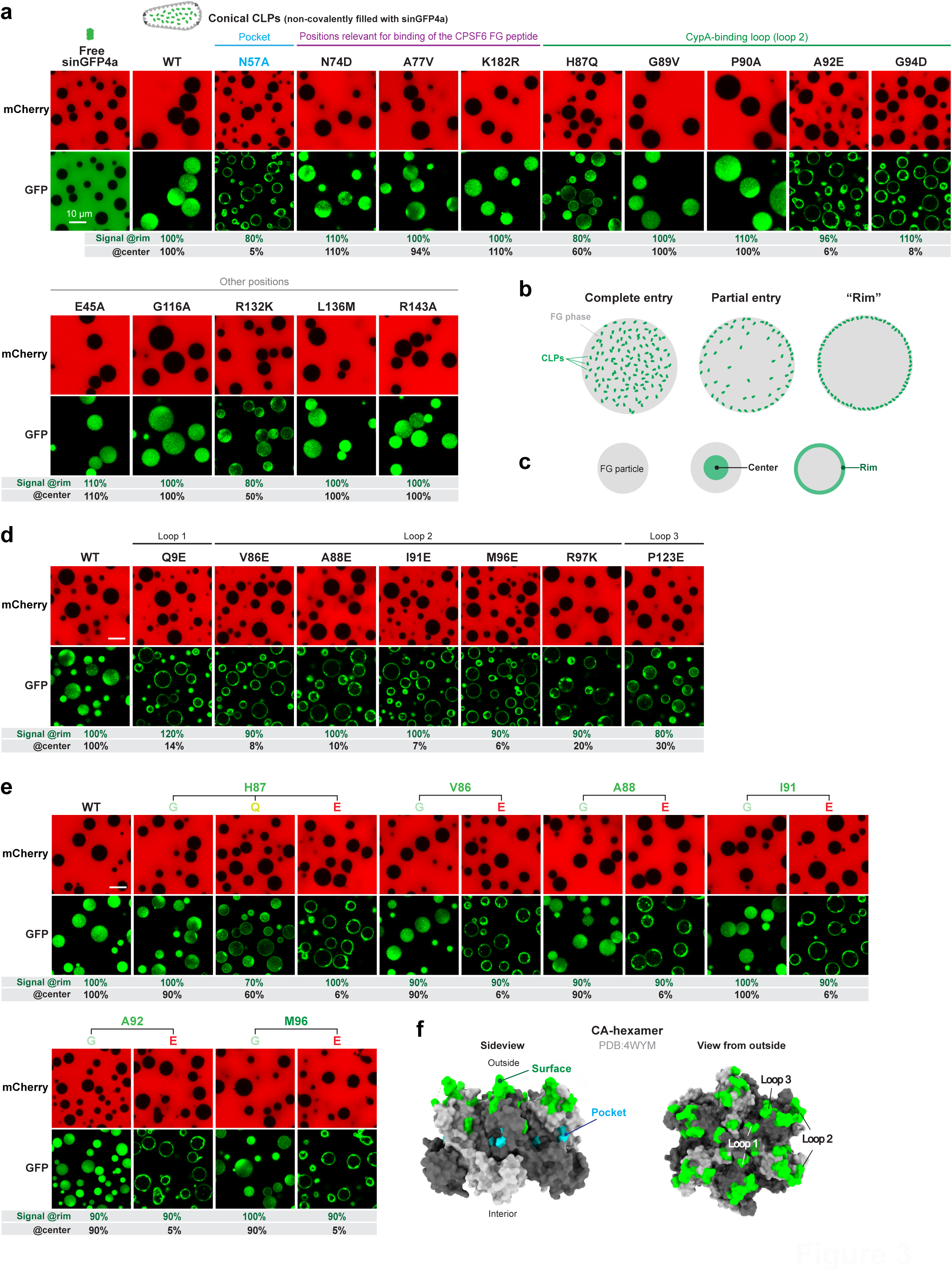
Identification of capsid mutations that impede FG phase-partitioning. Partitioning experiments into the FG phase (GLFG 12mer repeats) were performed as in Fig.1c, but using indicated (wildtype or mutant) CLPs non-covalently filled with sinGFP4a. Experiments were repeated independently with identical outcomes (*N* = 3). Scale bars, 10 μm. Negative stain electron micrographs, validating proper capsid assembly, are shown in Extended Data Fig.4. **a,** Comparison of wildtype CLPs and CLPs carrying previously reported mutations.Note the strong FG phase-partitioning defect of the N57A, A92E and G94D mutant CLPs. **b,** Illustration of the observed FG phase-partitioning phenotypes. **c,** Illustration of the quantification strategy. Fluorescence signals for the capsid species were integrated separately at the rim and in the center of FG particles, normalized to the wildtype values, and listed for each mutant in **a, d** and **e**. For more detailed quantifications, see Extended Data Fig.5. **d,** Drastic FG-partitioning defects in rationally designed capsid mutants, where FG-attractive residues had been exchanged for FG-repulsive ones (E or K). **e,** Selected CA positions were mutated to G, Q, or E, as indicated. Drastic FG phase-partitioning defects were evident only when an FG-repulsive glutamate was introduced. Mutations to an FG-neutral glycine or glutamine had mild effects at best. **f,** A CA-hexamer (PDB 4WYM) viewed from the side and the capsid’s outside. Surface mutant positions with FG-partitioning defects are colored in green, the N57 FG-binding pocket in cyan.

**Fig. 4.**
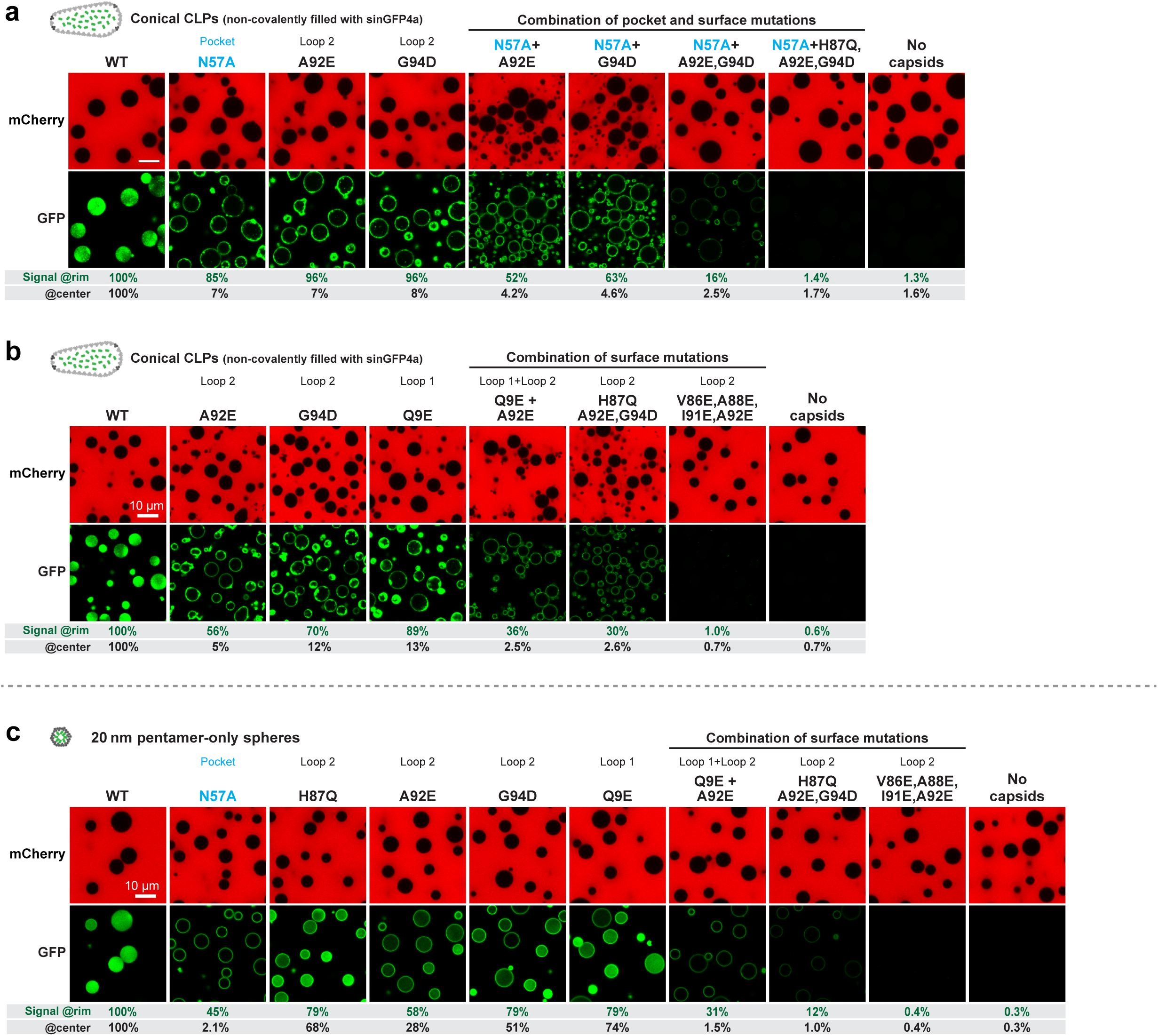
Synergy between ‘loss-of-FG-partitioning’ mutations in hexamer-dominated CLPs and pentamer-only spheres. **a, b,** Assay was performed as in Fig.3, analyzing WT CLPs and indicated mutants. Individual mutations caused already strong FG phase-partitioning defects. Combining mutations aggravated the defect up to a complete loss of partitioning. **c,** The assay was performed as in **a** and **b**, the difference being that pentamer-only spheres with a covalent EGFP-label were analyzed. These showed a very similar response to the mutations as the CLPs. Experiments were repeated independently with identical outcomes (*N* = 3). More detailed quantifications are shown in Extended Data Fig.7. Scale bars, 10 μm.

Following cytoplasmic microinjection into CPSF6-supplemented mouse oocytes, large CLPs inserted into NPCs, completed passage, and accumulated at nuclear speckles (Fig.6). With just endogenous CPSF6 levels, however, they remained arrested at NPCs. This can be explained by a lower CPSF6 availability in late-stage oocytes than in typical HIV target cells. At non-limiting CPSF6 levels, the capsid passage through NPCs was comparably efficient - considering that oocytes were loaded with far more capsids (∼10 000) than during a genuine infection and that images were acquired already 30 minutes post-injection, which is fast for a transport experiment in oocytes, where diffusion distances and thus diffusion times far exceed those in somatic cells. It is also fast compared to the estimated 1.5-hour capsid dwell time at NPCs during a genuine infection^105^. The kinetics of NPC-passage and release of these large capsids (Fig.6) thus appear to be within a physiologically plausible range.

**Fig. 5.**
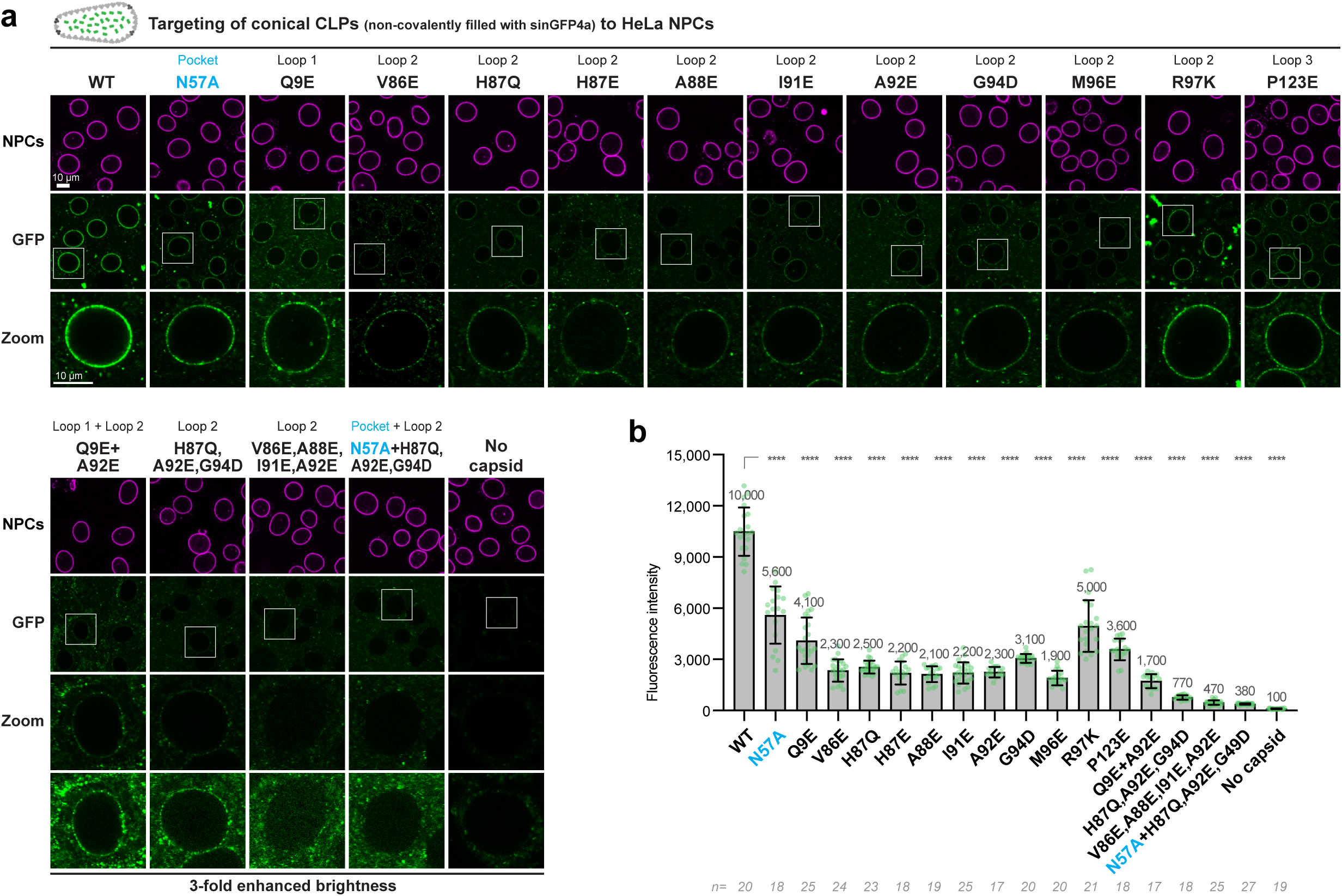
NPC-binding requires a lack of FG-repulsive residues on the capsid surface. **a,** Targeting of conical CLPs (non-covalently filled with sinGFP4) to HeLa cell NPCs was as in Fig.1e. Indicated mutants were tested with identical scan settings as for wild-type (WT) capsids. **b,** Quantification of GFP signals on NPCs with larger datasets. Numbers are means; bars indicate mean ± s.d.; n = number of quantified nuclei. *P*-values were calculated by one-way ANOVA with Tukey’s post hoc test. *P-values*: ****<0.0001. Experiments were repeated independently with identical outcomes (*N* = 3). Scale bar, 10 μm.

**Fig. 6.**
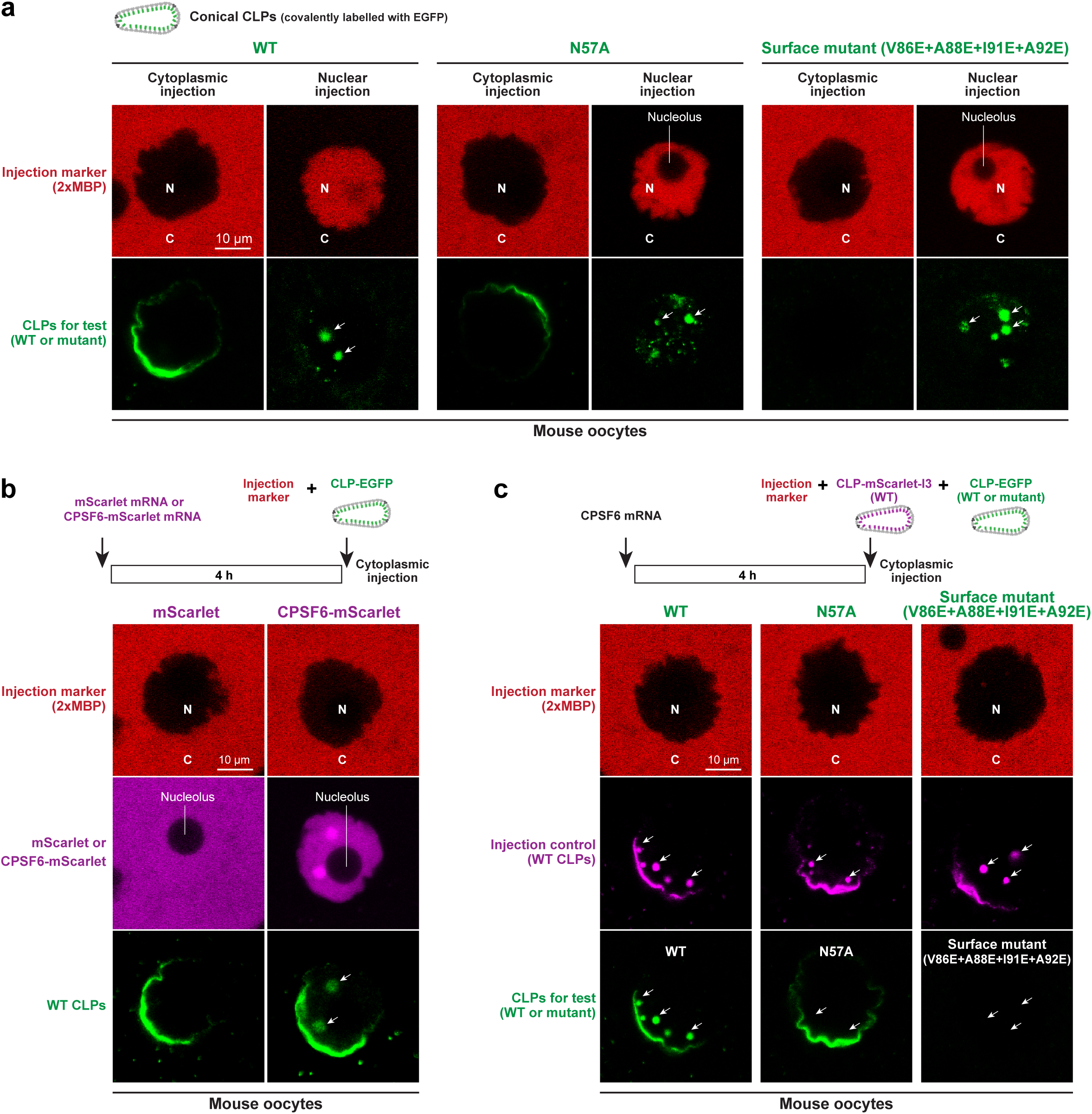
Requirements for NPC-passage of CLPs. **a,** Conical CLPs, carrying a C-terminal EGFP label and indicated mutations, were microinjected into fully grown mouse oocytes, and imaged 30 min later. The injected compartment, nucleus (N) or cytoplasm (C), was marked by co-injected Alexa647-labelled MBP dimers that do not cross the nuclear envelope (NE). Wildtype, or N57A, or V86E+A88E+I91E+A92 quadruple surface mutant CLPs accumulated in nuclear speckles after nuclear microinjection. This is a crucial control for the experiment of panel **c**. None of the large CLPs reached the speckles when microinjected into the cytoplasm. WT and N57A CLPs accumulated at the NE, the quadruple surface mutant did not – consistent with its failure to partition into an FG phase (see Fig.4b). The uneven accumulation of cytoplasmically injected capsids at the NE is due to the essentially irreversible binding upon first encounter of NPCs, combined with slow diffusion of the very large capsids and the resulting concentration gradient in the cytoplasm. **b,** mScarlet or a CPSF6-mScarlet fusion were over-expressed from mRNA that was microinjected into mouse oocytes. After four hours, EGFP-labelled wildtype CLPs were microinjected into the cytoplasm and imaged 30 minutes later. The CPSF6 fusion allowed the CLPs to pass NPCs and to accumulate in speckles, where they co-localized with CPSF6. **c,** Unlabelled CPSF6 was overexpressed from microinjected mRNA for four hours. Then, mScarlet-I3-labelled wildtype CLPs were co-injected with indicated EGFP-labelled mutant CLPs into the cytoplasm, and imaged 30 minutes later. The mScarlet-I3-labelled wildtype CLPs bound NPCs, crossed the NE, and accumulated in speckles. EGFP-labelled wildtype CLPs behaved the same. N57A mutant CLPs bound to the NE, but failed to pass. Quadruple surface mutant CLPs failed to bind NPCs/ the NE and failed to accumulate in speckles – again consistent with their failure to partition into an FG phase. Experiments were repeated independently with identical outcomes (*N* = 3). Scale bars, 10 μm.

To ‘extract’ the capsid from its highly multivalent interactions with the FG phase, CPSF6 must compete against an extremely high local FG concentration. Given this challenge, CPSF6’s block of capsid-partitioning is remarkably strong – far stronger than that of the N57A mutation, which essentially inactivates the FG-binding pocket (Fig.7). This suggests that CPSF6 not only obstructs the pocket but also masks the FG-attractive surface identified here.

**Fig. 7.**
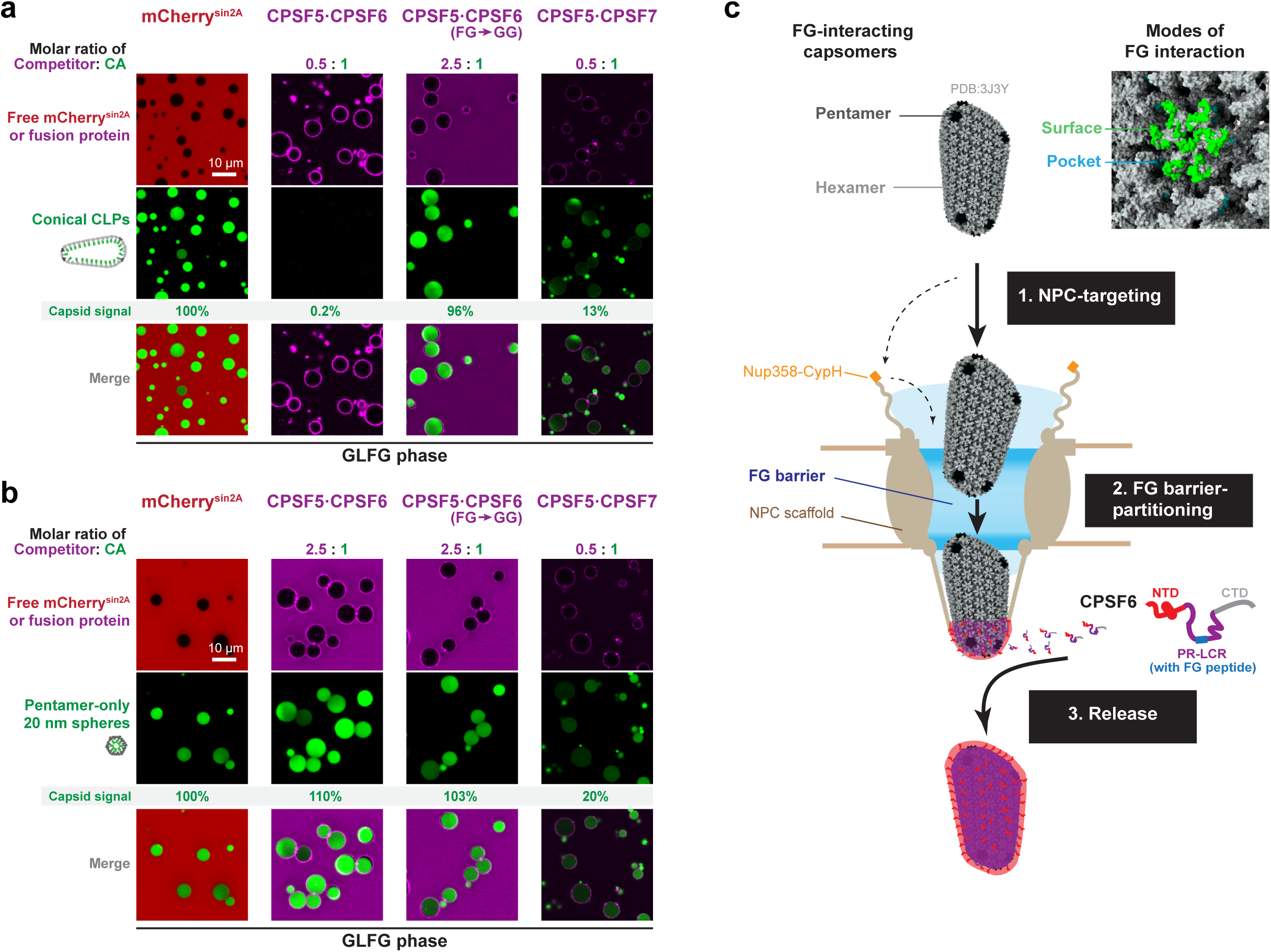
CPSF6 potently antagonizes capsid-partitioning into an FG phase. **a,** Capsid-partitioning into an FG phase was assayed as in Fig.3, but before addition to the GLFG phase, CLPs were pre-mixed with indicated protein complexes. The CPSF5·CPSF6 complex blocked FG-partitioning completely, the CPSF6^F284G^ mutant complex had no effect. CPFS7 (a CPSF6 paralog), which also contains a singular FG motif, competed the partioning moderately. Molar ratios were calculated for the tetrameric (A_2_B_2_) CPSF5·CPSF6 or CPSF5·CPSF7 complexes. **b,** Experiment was as in panel **a**, but FG phase-partitioning of 20 nm pentamer-only spheres was analyzed. Note that this capsid species partitioned in a CPSF6-resistant manner. This documents that the re-arranged pentamer N57 pocket has no defect in binding ‘normal’ FG peptides but a severe defect in binding the sterically constrained CPSF6 FG peptide. CPSF7 inhibited the partitioning moderately, and thus binds the pocket in a more tolerant manner than CPSF6 does. Fluorescence signals for the capsid species were measured in the center of FG particles, normalized to the wildtype values. Experiments were repeated independently with identical outcomes (*N* = 3). Scale bars, 10 μm. **c,** Scheme of HIV-1 capsid passage through NPCs. To reach the nucleus, the HIV capsid has to dock to NPCs first. The initial interaction might be a capsid-CypH interaction or a direct targeting to the FG barrier. The capsid then fully partitions into the FG phase, which probably requires disengagement from the CypH domain. CPSF6 extracts the capsid from the NPC barrier and promotes release into the nucleus, by using a competing FG peptide and forming a condensate around the capsid, which not only provides a very high local CPSF6 concentration but also masks the FG-attractive surface of the capsid and provides an FG-repulsive outer layer instead. The capsid is thus handed over from one phase to another. See main text and Suppl. Figure 3 for a more detailed description.

Surface masking probably involves fuzzy hydrophobic interactions with the proline-rich low-complexity region (PR-LCR) that flanks the CPSF6 FG peptide and contributes to capsid-binding^106^. The PR-LCR is extremely depleted of charged FG-repulsive residues but enriched in hydrophobic ones (Supplementary Fig. 3), resembling an Nup98 FG domain. Thus, just as the capsid is attracted to the FG phase, the PR-LCR is attracted to the compositionally biased capsid surface. This attraction allows capsids to seed CPSF6 condensates^88,88,107^, which then provide a very high local CPSF6 concentration and thus avidity to the interaction.

An intriguing twist is the negatively charged CPSF6 N-terminus, which appears to be FG-repulsive and immiscible with the local PR-LCR condensate. This suggests a layered arrangement around the capsid: The PR-LCR condensate forms the inner layer, masks the FG-attractive surface, blocks the FG-pocket (via the embedded FG motif), and provides condensate-stabilizing interactions (perhaps together with the C-terminal R-rich domain). The negatively charged (FG-repulsive) CPSF6 N-terminus is excluded from the condensate and thus forms an FG-repulsive outer layer.

This arrangement would thus cause the capsid to switch from FG-attractive to FG-repulsive, explaining capid release from the FG phase of NPCs. The nuclear localization of CPSF6 ensures that this release occurs into the nucleus and makes the capsid transport directional. In this respect, CPSF6 appears to be analogous to RanGTP, which also acts as a directional switch, namely as a nucleus-specific release factor for cellular cargoes from importins and thus from NPCs^4^.

## Methods

### DNA sequences for recombinant protein expression

All recombinant proteins used in this study were produced in *E.coli*, using codon-optimized genes, His_14_-NEDD8 or His_14_-SUMO tags and a purification strategy that includes binding to a Ni^2+^ chelate matrix and proteolytic release by NEDP1 or SenP1/ Ulp1 (ref.^110^). Expression vectors are listed in Supplementary Tables S2-S5.

### Assembly and purification of conical CLPs

His_14_-bdSUMO tagged CA^P1A^ was expressed in NEB Express *E. coli* cells (New England Biolabs, C2523). Here, the P1A mutation is required for tag-removal, i.e. to allow SUMO cleavage. Induction was with 0.1 mM IPTG (isopropyl-β-thiogalactoside) at 18°C for 16 hours. Cells were harvested by centrifugation, resuspended in lysis buffer (50 mM Tris/HCl pH 8.0, 300 mM NaCl, 20 mM imidazole, 1 mM TCEP), and lysed by a freeze-thaw cycle followed by sonication. The lysates were cleared by ultracentrifugation, the soluble fractions were bound to Ni^2+^ chelate beads for 1 hr at 4 °C, beads were washed with wash buffer (50 mM Tris/HCl pH 8.0, 40 mM imidazole, 300 mM NaCl, 1 mM TCEP), and proteins eluted by tag cleavage with 100 nM of bdSENP1 protease in cleavage buffer (50 mM Tris/HCl pH 8.0, 20 mM imidazole, 300 mM NaCl, 0.5 mM TCEP) for 3 hours at 4 °C.

The tag-free proteins were concentrated to approximately 20-30 mg/ml. IP6 (D-myo-inositol hexakisphosphate)-assisted assembly^64^ into CLPs essentially followed a published protocol^73^ with minor modifications. The buffer of the CA protein was exchanged to 25 mM MES pH 6.0, 50 mM NaCl and 1 mM TECP. Assembly was initiated by adding 0.5 volumes of 75 mM MES pH 6.0, 150 mM NaCl, 15 mM IP6, 3 mM TCEP and shifting the temperature to 37°C for two hours. The final volume was 500µl and the CA concentration was 12 mg/ml. We generated more homogenous CLPs (size from ∼60×100 nm to ∼80×160 nm) with this protocol than with the previous one^33^, which produced larger particles in a different assembly buffer (50 mM Tris pH 8.0, 1M NaCl and 0.1mM IP6).

Assembled CLPs were pelleted by centrifugation in an 5424R Eppendorf microcentrifuge (FA-45-24-11 rotor) at 21 000 g for 10 min, resuspended in gel filtration buffer (25 mM Tris/HCl pH 8.0, 150 mM NaCl, 0.5 mM IP6 and 0.5 mM TCEP), aggregates were removed by centrifugation @ 21 000 g for 5 minutes. Note that the pH-shift prevents the pelleting of properly assembled CLPs in the second centrifugation step. The CLPs in the supernatant were further purified by size exclusion chromatography on a Superose6 Increase 10/300 GL column (equilibrated in gel filtration buffer), where they eluted in the void volume. Assembly was performed either in the presence of 2 mM soluble sinGFP4a (ref.^14^), as a non-covalently encapsulated tracer, or with CA-EGFP or CA-mScarlet-I3 fusion (ref.^111^) used as a tracer in a 1:6 molar ratio to the unlabeled CA.

CLPs with the Q9E-A92E, H87Q-A92E-G94D, N57A-A92E-G94D, N57A-H87Q-A92E-G94D and V86E-A88E-I91E-A92E mutations do not pellet in the post-assembly centrifugation step, probably because of charge-repulsion. They were therefore directly applied to the Superose 6 column, after removing aggregates by a 5 minutes 21 000 g centrifugation step.

### Assembly and purification of 40 nm capsid spheres

His_14_-bdSUMO tagged CA^P1A^ with the additional N21C, A22C mutations^76^ was expressed and purified as described above and concentrated to approximately 6 mg/ml. Assembly was performed by dialysis against 50 mM Tris/HCl pH 8.0, 1 M NaCl and 0.1 mM IP6 for 24 to 48 hours. Assembled CLPs were further purified by size exclusion chromatography on a Superose6 Increase 10/300 GL column equilibrated in 25 mM Tris/HCl pH 8.0, 500 mM NaCl, 0.5 mM IP6, where they eluted near the void volume.

### Assembly and purification of 20 nm pentamer-only spheres

For assembly of 20 nm pentamer-only spheres, His_14_-bdSUMO tagged CA^P1A,G60A,G61P^ was purified as described above and concentrated to approximately 20-30 mg/ml. Assembly into 20 nm capsids followed a published protocol^73^ with minor modifications. Briefly, the buffer of the CA protein was exchanged to 25 mM MES pH 6.0, 50 mM NaCl and 1 mM TECP, assembly was initiated by adding 0.5 volumes of 75 mM MES pH 6.0, 150 mM NaCl, 15 mM IP6, 3 mM TCEP, and allowed to proceed in a volume of 500 µl for two hours at 37°C and a CA concentration of 12 mg/ml. Assembled CLPs were further purified by size exclusion chromatography on a Superose6 Increase 10/300 GL column equilibrated in 25 mM Tris/HCl pH 8.0, 150 mM NaCl, 0.5 mM IP6 and 0.5 mM TCEP.

### Fluorescence labeling

The anti-Nup133 nanobody xhNup133-Nb2t was labelled with Alexa Fluor™ 647 C_2_ maleimide (ThermoFisher) through two ectopic cysteines at the N- and C-termini as desribed^79^, reaching a density of labeling (DOL) of 2. The MBP tandem dimer (2xMBP) was also labeled with Alexa Fluor™ 647 C_2_ maleimide but through a single cysteine at the C terminus.

### Negative stain electron microscopy

Samples were bound to a glow-discharged carbon foil covered 400 mesh copper grid. After successive washes with water, samples were stained with 1% uranyl acetate aq. and examined at room temperature on a Talos L120C transmission electron microscope (Thermo Fisher Scientific).

### Digitonin-permeabilized cell assays

HeLa K cells (RRID: CVCL_1922) and XTC-2 cells (RRID: CVCL_5610; ref.^81^) were obtained from the European Cell Culture Collection, authenticated by the manufacturer, and tested negative for mycoplasma. HeLa cells were grown at 37°C in Dulbecco’s modified Eagle’s Medium (DMEM, high glucose), supplemented with 10% heat-inactivated fetal calf serum (FCS), antibiotics (“AAS”, Sigma-Aldrich) and 5% CO_2_. XTC-2 cells were cultivated at 22°C in 70% Leibovitz medium (diluted with water), 10% FCS, AAS antibiotics.

Cells were seeded on 8-well μ-Slides (IBIDI, Germany) to 70% confluence. Plasma membranes were permeabilized^32,112^ by treating the cells with 30 μg/ml digitonin (water-soluble fraction) in transport buffer (20 mM HEPES/KOH pH 7.5, 110 mM (HeLa) or 80 mM potassium acetate (XTC-2), 5 mM magnesium acetate, 0.5 mM EGTA, 250 mM sucrose) for 3 or 6 mins at 25°C (with gentle shaking), followed by three washing steps in transport buffer. Permeabilized cells were then incubated for 30 minutes with 40 nM Alexa647-labeled xhNup133-Nb2t nanobody and either EGFP (3 μM), CA-EGFP (1 μM), conical CLPs, 40 nm capsid spheres of HIV-1 or 20 nm pentamer-only spheres (with CA concentrations of 5, 1, and 2 μM, respectively). The samples were then directly scanned with a Leica SP8 confocal laser-scanning microscope (equipped with a 63x oil objective and HyD GaAsP detectors), with sequential excitation at 488 and 638 nm.

### mRNA for microinjection

Mouse CPSF6 (CCDS Database: CCDS78898.1) and mScarlet^113^ cDNAs were cloned into pGEMHE^114^, which contains a T7 promoter, *Xenopus* globin 5’ and 3’UTRs, and poly(A) tail. Plasmids were linearized by AscI (NEB) before *in vitro* transcription using the HiScribe T7 ARCA mRNA Kit (NEB #E2060S). mRNAs were purified with the RNeasy Mini Kit (Qiagen #74104). 4 pl of 1 µM mRNA was cytoplasmically injected.

### Microinjections

Mouse oocytes were obtained from ovaries of 9-week-old CD1 mice that were maintained in a specific pathogen-free environment according to the Federation of European Laboratory Animal Science Association guidelines and recommendations, as previously described^115^. Fully-grown oocytes were kept arrested in prophase in homemade M2 medium supplemented with 250 μM dibutyryl cyclic AMP under paraffin oil (NidaCon) at 37°C. Labelled CLPs, along with the Alexa647 labelled 2xMBP injection marker, were microinjected into cytosol or nucleus of oocytes, as previously described^33^. Oocytes were imaged about 30 minutes after microinjection.

### FG phase assays

The assays were performed as previously described^78^ with minor modifications. In brief, 1mM FG domain stocks were prepared in 2 M (GLFG repeats) or 4 M guanidinium hydrochloride (all other repeat domains). Phase separation was initiated by rapid dilution of the FG domain stock with 25 volumes (GLFG repeats) or 50 volumes (other repeats) of assay buffer (50 mM Tris/HCl pH 7.5, 150 or 250 mM NaCl, 0.5 mM IP6), followed by a further fourfold dilution in buffer with indicated fluorescent probes. The resulting mixture was pipetted on collagen-coated μ-slides 18-well (IBIDI, Germany), and FG particles were allowed to settle on the bottom for one hour, before confocal scans were taken. Salt effects of the assay are detailed in Extended Data Fig.10.

Partition coefficients were calculated as the integrated raw signal within independent FG particles (In) divided by the signals reference areas in outside regions (Out). Background was not subtracted, which means that the numbers of high partition coefficients are still underestimated. Plots are shown for representative FG particles (with 5-10 μm diameters). Images were analyzed in FIJI 2.9.0 and the exported data were further processed in GraphPad Prism 9.5.1.

The sequences of FG domains used in FG phase assays are shown in Supplementary Table S1.

### Structure modeling

The Alphafold 3 Server^116^ was used for all modellings. FG-binding was modelled with 6 copies of the wildtype HIV-1:M CA and 6 copies of the respective FG peptide. The following sequences were used: CPSF6 (PVLFPGQPFGQPPLG), GLFG (QPATGGLFGGNTQ), SLFG (QPATGSLFGGNTQ), FSFG (NTQPATGFSFGGNTQPATG)

CA (PIVQNLQGQMVHQAISPRTLNAWVKVVEEKAFSPEVIPMFSALSEGATPQDLNTMLNTVGGHQAAMQMLKETINEEAAEWDRLHPVHAGPIAPGQMREPRGSDIAGTTSTLQEQIGWMTHNPPIPVGEIYKRWIILGLNKIVRMYSPTSILDIRQGPKEPFRDYVDRFYKTLRAEQASQEVKNWMTETLLVQNANPDCKTILKALGPGATLEEMMTACQGVGGPGHKARVL)

For the structural alignment and assessment of accessibilities (Extended Data Fig.9), the sequences shown in the figure were used as inputs to model mature hexamers.

---

**Extended Data Fig. 1.**
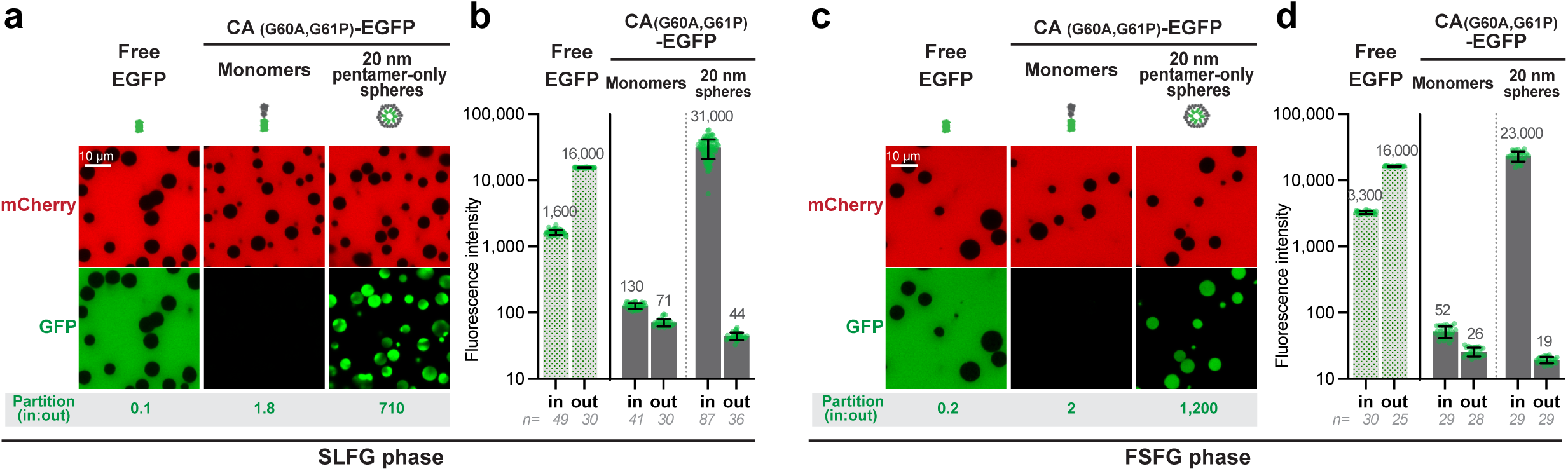
Pentamer-only capsids partition efficiently into SLFG and FSFG phases. FG phase-partitioning experiments and quantifications were performed as in Fig.1c-d, but FG phases were assembled from different FG repeat domains. Note that fully assembled T1 spheres partitioned very well also in these alternative FG phases – 500 times better than the unassembled CA monomers. Scan settings were identical for each monomer-capsid pair. Experiments were repeated independently with identical outcomes (*N* = 3). Scale bars, 10 μm. **a,b,** Instead of the 52x GLFG 12mer repeats (G**GLFG**GNTQPAT)_52_ as in Fig.1, a domain with 52 SLFG 12mer repeat units was used (ref.^31^; see Supplementary Table S1 for sequence). The inter-FG spacer was also free of charged amino acids – a typical feature of Nup98-type FG repeats. SLFG is the second most abundant FG motif in Nup98 FG repeat domains. **c,d,** An FG repeat domain of 669 residues with non-charged spacers and 36 FSFG motifs was used. The lower motif density (1 motif per 18 residues) considers that FSFG motifs are more hydrophobic and thus more cohesive than GLFG or SLFG motifs. For sequence see ref.^31^ or Supplementary Table S1.

**Extended Data Fig. 2.**
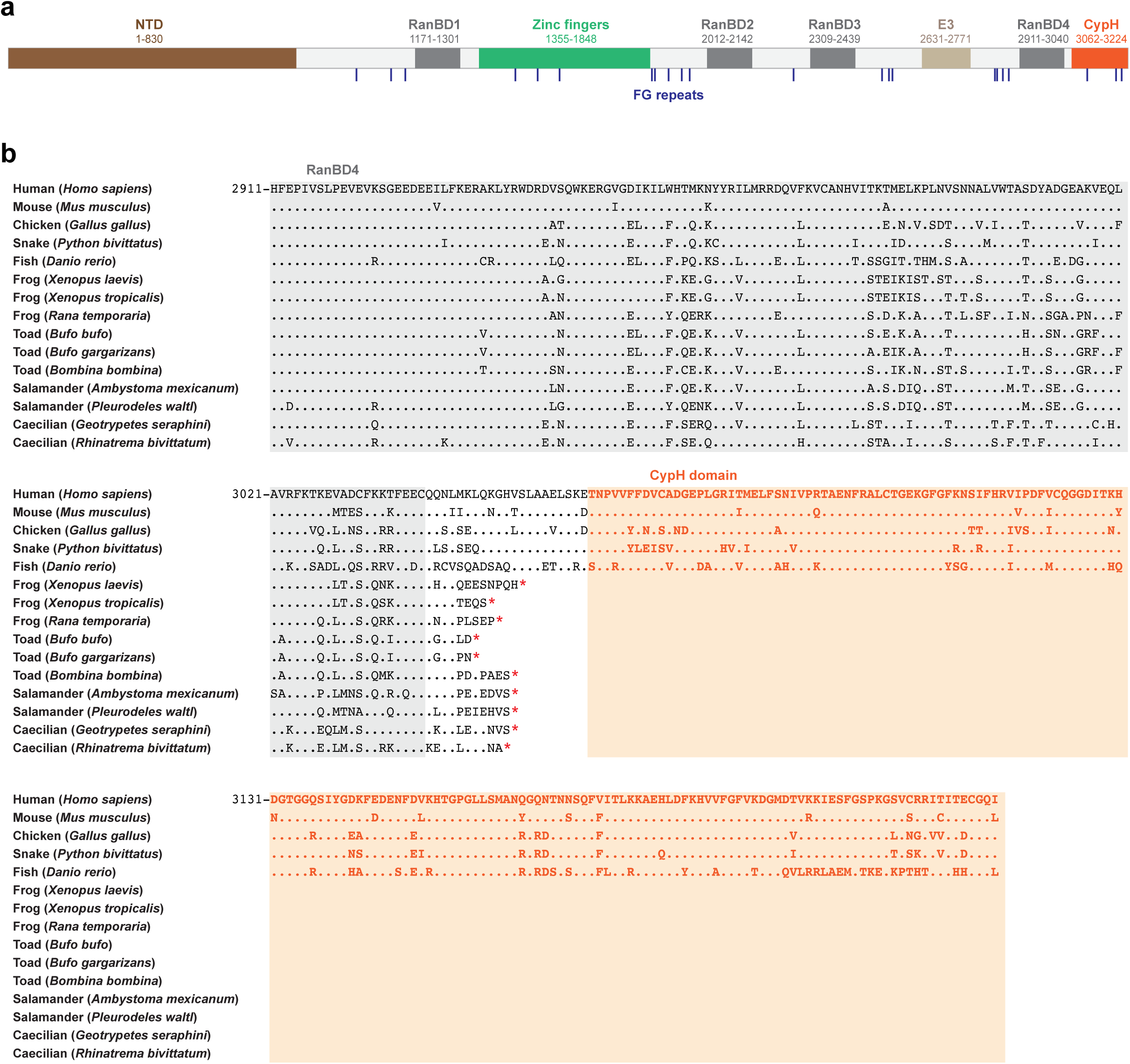
*Xenopus* Nup358/ RanBP2 lacks the C-terminal CypH (cyclophilin homology) domain. **a,** Domain organization of human Nup358, which includes an N-terminal NPC-anchor domain, four RanBP1-like Ran-binding domains (RanBD1-4), 8 Ran-binding zinc finger domains, two SUMO-E3 ligase domains, several FG repeat domains, and a C-terminal CypH domain. **b,** Sequence alignment of the C terminus part of Nup358 from indicated species. Note that Nup358 from mammals, birds, reptiles, or fish all have a C-terminal CypH domain. However, the CypH domain has been lost in modern amphibians such as frogs, toads, salamanders, and caecilians, where a translation stop is found immediately after the last RanBD. Cells from the frog/ toad *Xenopus laevis* can therefore be considered as a naturally occurring ΔCypH mutant. Sequence identifiers are listed in Supplementary Table S6.

**Extended Data Fig. 3.**
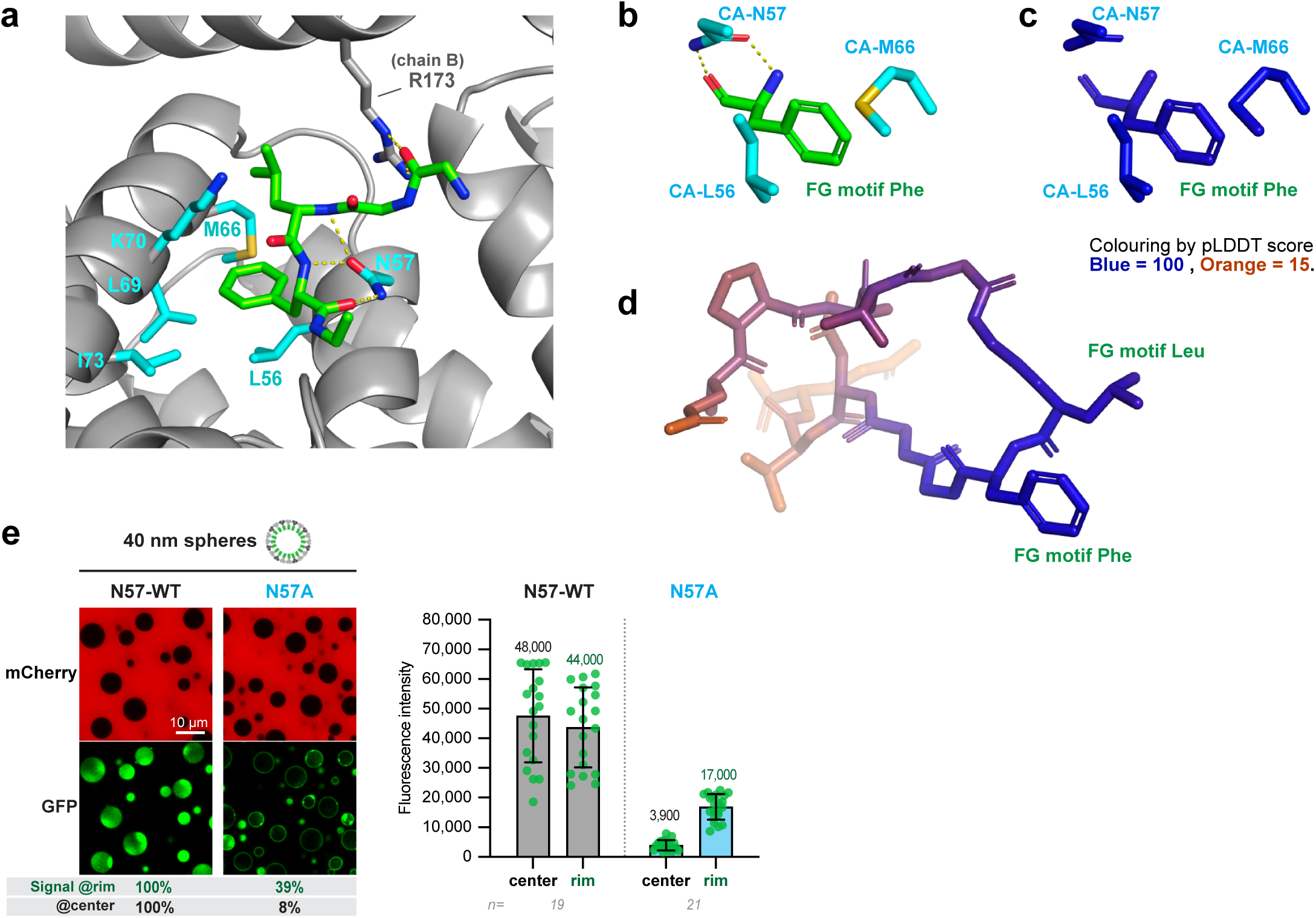
Alphafold 3-modeling confirms that the N57 pocket binds diverse FG repeats in a similar hydrogen bond-stabilized manner. **a,** A mature full-length CA-hexamer was modeled with GLFG peptides bound (the same repeats as in the FG phase assays of figE. 1, 3-4). Images show just the two CA chains (chain A and chain B) that accommodate the corresponding peptide. The N57 sidechain amide forms two hydrogen-bonds to the backbone of the FG phenylalanine – virtually identically to the FG motif of the CPSF6 peptide (PDB 4B4N, ref.^56^). The N57 sidechain oxygen forms a third H-bond to the backbone nitrogen of the preceding leucine. Additional H-bonds are formed between the guanidine group of R173 (chain B) and the backbone carbonyl oxygen of the glycine or serine that precedes the LFG motif. The sidechain of each FG phenylalanine engages in extensive hydrophobic contact with L56, M66, L69, (the aliphatic part of) K70, and I73 of chain A. CA-hexamer modeling with SLFG and FSFG peptides are shown in Supplementary Figure S1. **b,** The same highest-ranking structure of a GLFG peptide bound as in (**a**), but panel shows just the phenylalanine of the FG motif, which mediates the key interactions, namely its backbone H-bonding to the sidechain of N57 and the phenyl moiety engaging in hydrophobic contacts with M66 and L56. Coloring is by chain and elements. **c,** Same view as **b** but colored by the local pLDDT score, using the ‘spectrum b’ function of Pymol with indicated settings. The very high score (blue color) indicates a very high confidence of this part of the model (pLDDT= 94). **d,** The entire FG peptide in pLDDT coloring; CA residues are omitted. Note that the pLDDT score deteriorates with sequence distance from the FG motif, reflecting flexibility in accommodating diverse FG repeats. **e,** Wildtype and N57A mutant 40 nm spheres were allowed to partition into GLFG phase, essentially as described in Fig.1 and ref.^33^. The N57A mutations caused clear partitioning defects, documenting that the N57 hydrogen bonds contribute to relevant FG interactions.

**Extended Data Fig. 4.**
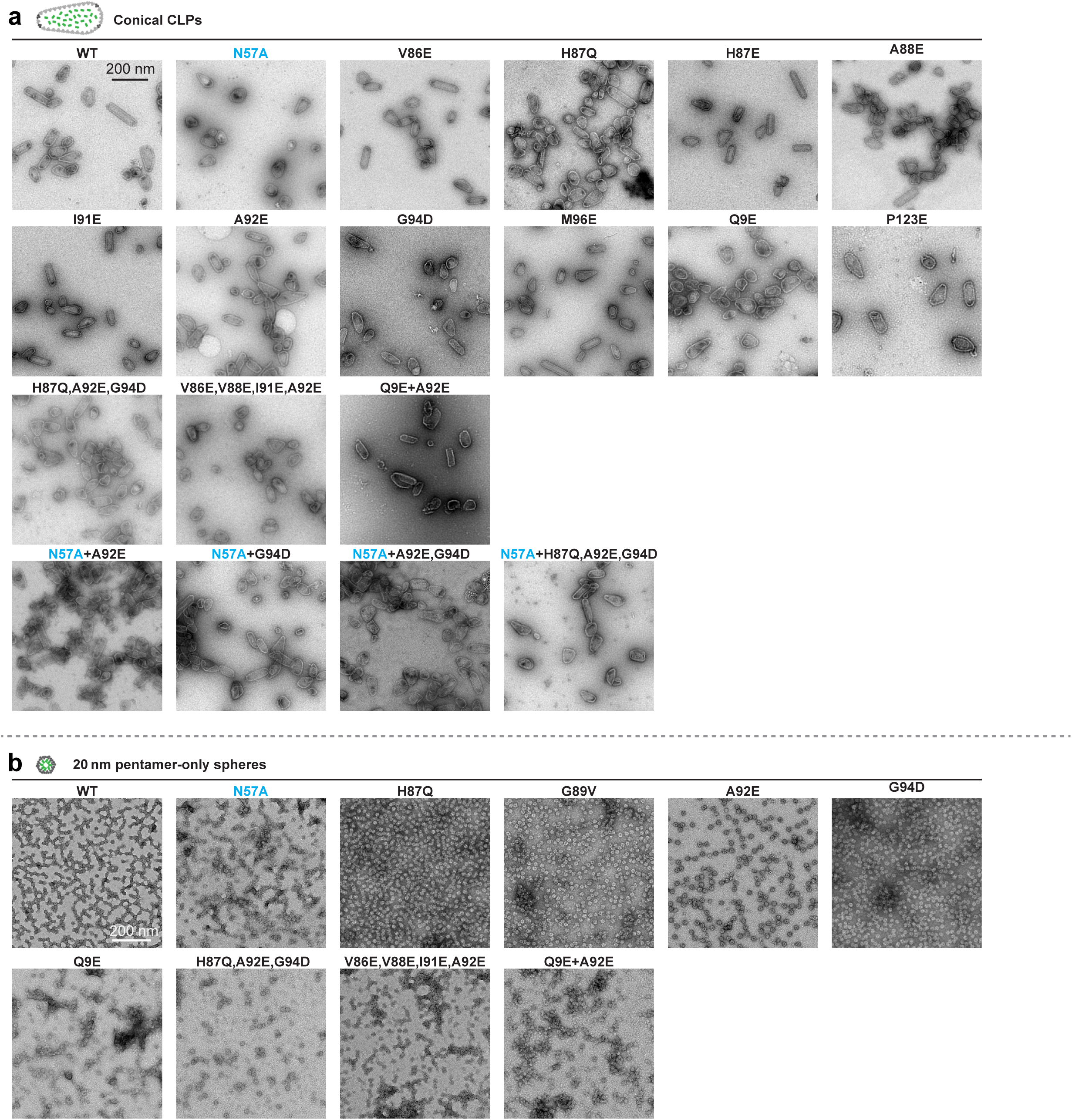
Negative stain electron micrographs of mutant HIV-1 capsids. **a,** Wildtype or mutant CA was recombinantly expressed, purified, and allowed to assemble into cone-shaped CLPs (in the presence of IP6; see Methods). The assemblies were applied to a Superose 6 size exclusion column, where CLPs elute close to the void volume. This can be considered as a first criterion for a faithful CLP assembly. The corresponding fractions were pooled and further analyzed by negative stain electron microscopy. Representative images are shown in this panel and confirm the proper assembly of all mutants for which we describe a transport phenotype. Several other mutants showed defects in capsid assembly (for example, Q7E, M10E/K, V11E, P122E, I124E). These were excluded from the analysis. **b,** Wildtype and mutant pentamer-only spheres were assembled as analyzed in **a**, the differences being (1) that the CA protein contained in addition the G60A,G61P double mutation, which is required to assemble pentamer-only spheres^73^, (2) that the assemblies were smaller than the CLPs (20 nm instead of 100-200 nm), and (3) that these smaller assemblies also eluted later from the Superose 6 column. Scale bars, 200 nm.

**Extended Data Fig. 5.**
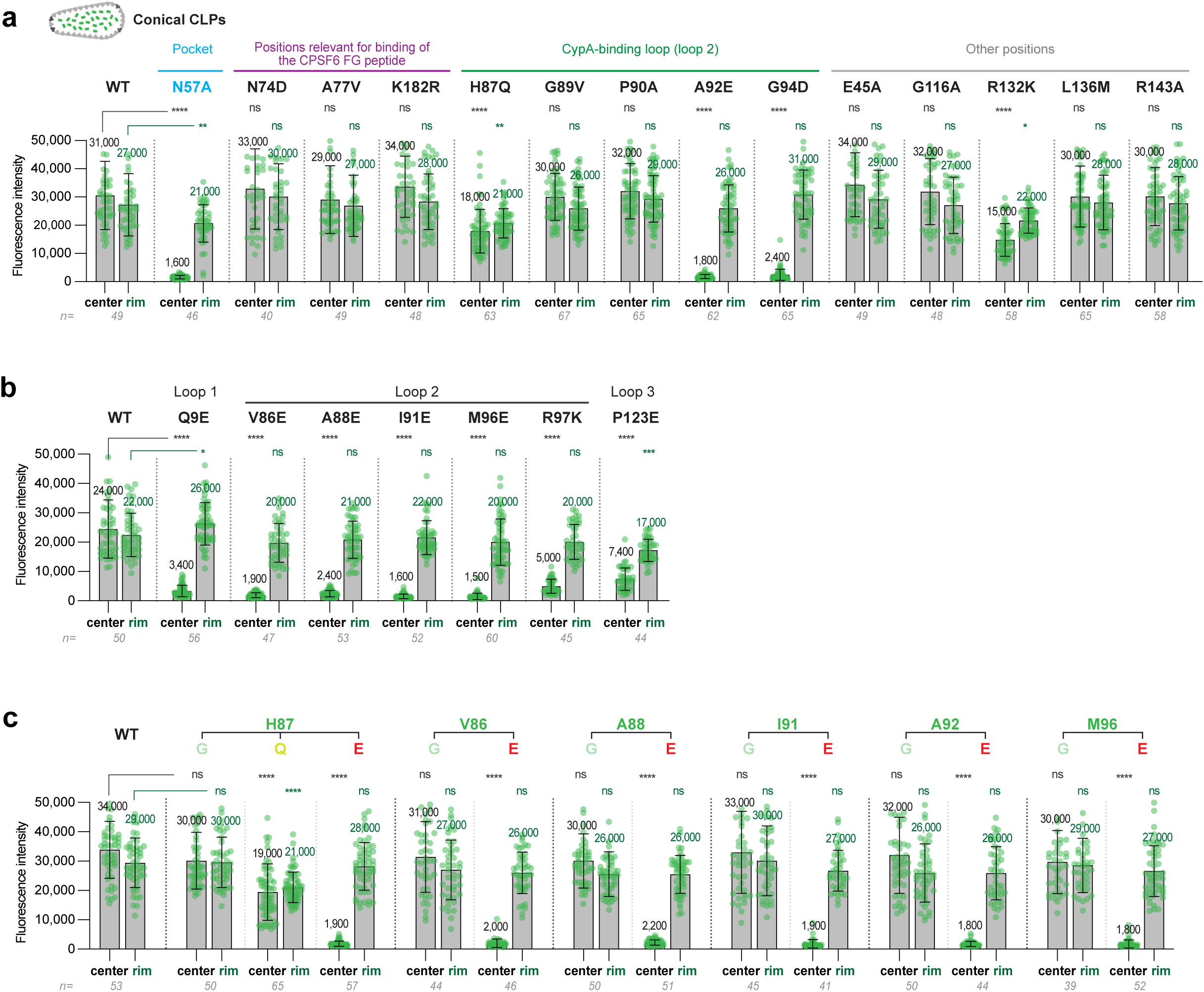
Identification of capsid mutations that caused defects in FG phase-partitioning. The panels provide more detailed quantifications for the partitioning of capsid mutants into the GLFG phase of Fig.3. They are based on larger datasets. Capsid signals at the FG phase surface and in the FG phase center were integrated separately. *P*-values were calculated using one-way ANOVA with Tukey’s post hoc test. *P-values*: * < 0.05, ** < 0.01, *** < 0.001, ****<0.0001, ns: not significant (*P-value* ≥0.05).

**Extended Data Fig. 6.**
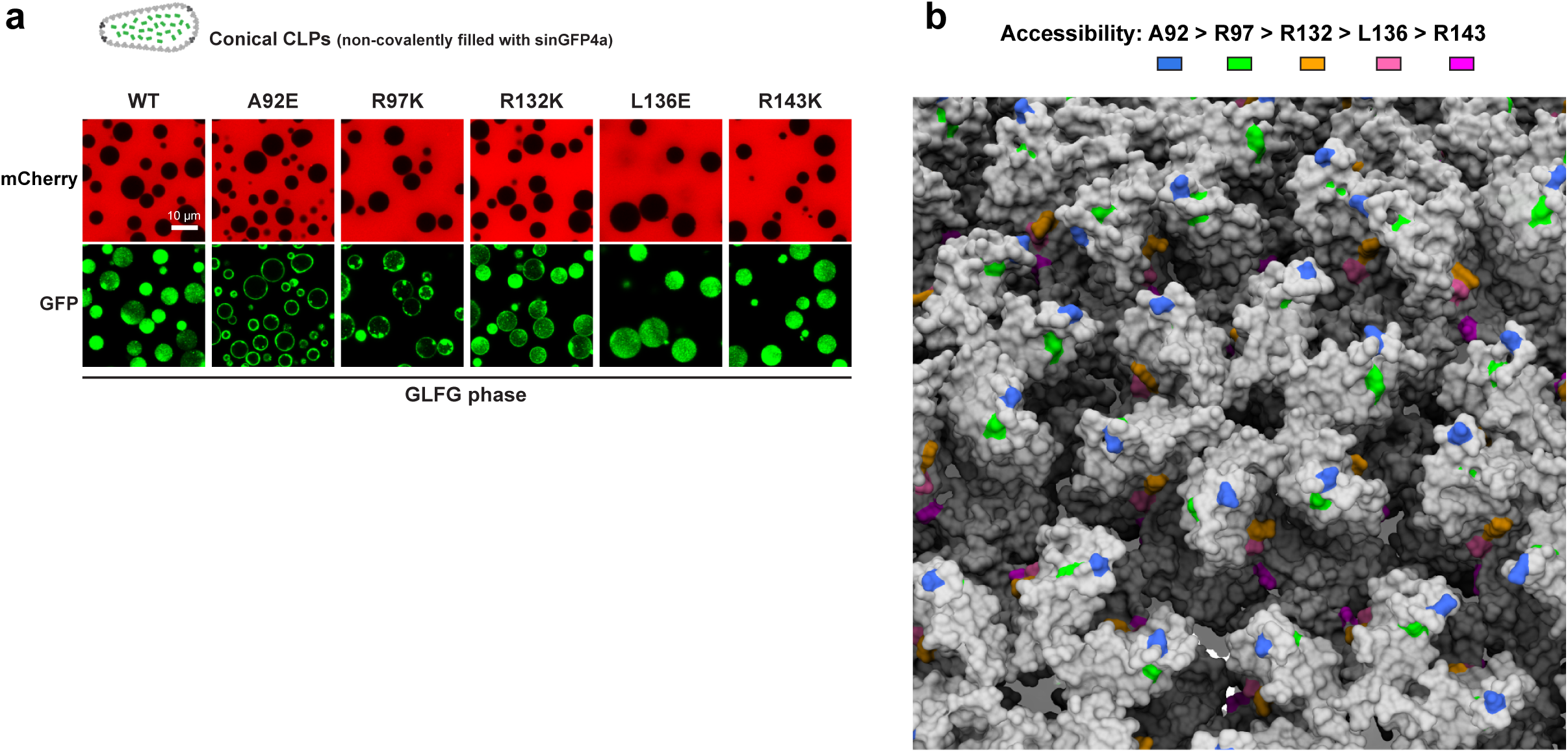
Reduction of FG phase-solubility by FG-repulsive capsid mutations depends on the accessibility of the mutated residues. **a,** The FG phase-partitioning of CLPs was tested as in Fig.3, using wild-type and indicated mutants. Partitioning was strongly inhibited by FG-repulsive mutations at highly accessible residues (A92E and R97K). When similar mutations were introduced at less accessible positions, partitioning was little affected (R132K) or unaffected (L136E or R143K). **b,** A92, R97, R132, L136 and R143 are shown with indicated coloring in a capsid structure (PDB 3J3Y).

**Extended Data Fig. 7.**
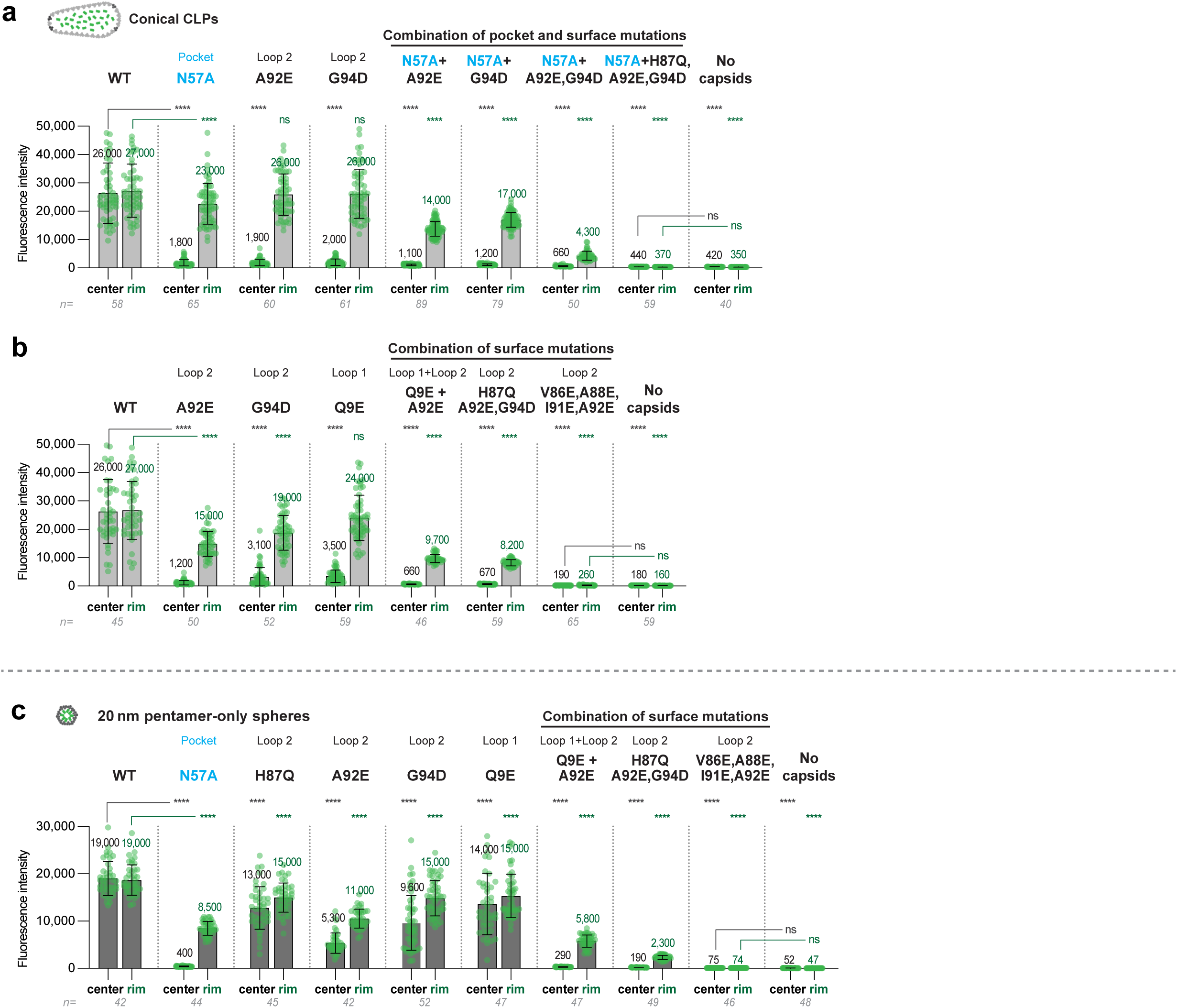
Synergy between surface residues and the FG-binding pocket in FG phase-partitioning of the capsid. Quantification of the capsid-partitioning into the GLFG phase as shown in Fig.4 but based on larger datasets. Quantification was performed as in Extended Data Fig.5. *P-values*: ****<0.0001, ns: not significant (*P-value* ≥0.05).

**Extended Data Fig. 8.**
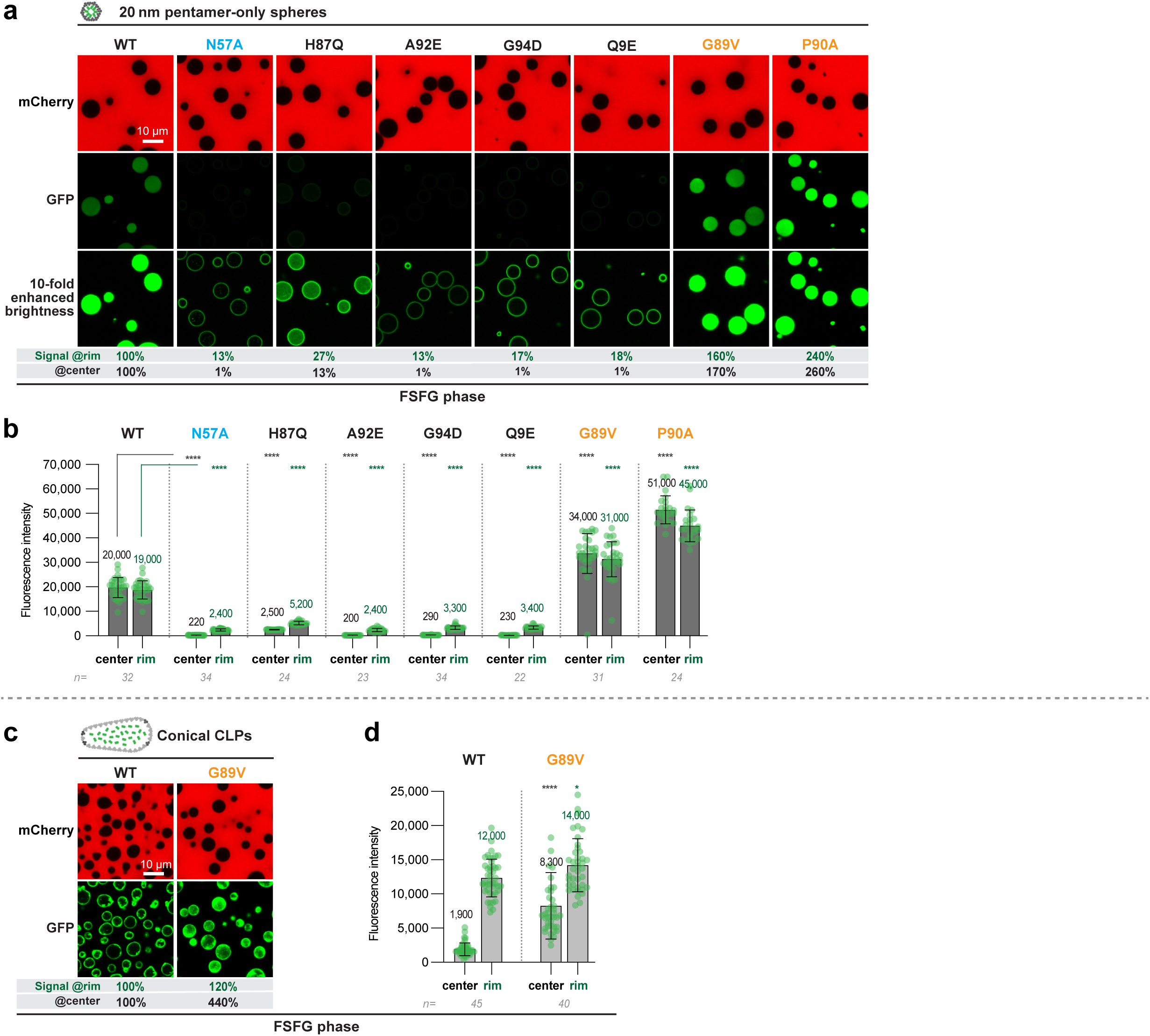
Mutations that block or enhance the capsid-partitioning into an FSFG phase a,b, Partitioning of pentamer-only spheres into the FSFG phase was as in Extended Data Fig.1, but the indicated CA mutants were also analyzed. The resulting partitioning defects by N57A, A92E, H87Q, G94D, and Q9E exchanges were considerably more severe as compared to the GLFG phase, showing an up to 100-fold reduction of WT. This can be explained by the stronger cohesion of the more hydrophobic FSFG motifs, which necessitates stronger capsid-FG interactions for melting into the phase. Interestingly, the experiment also revealed that the G89V and P90A mutations enhance the partitioning of this capsid species. The effects can be explained by G89V making the surface more hydrophobic and by the P90A exchange allowing for an additional hydrogen-bonding between the nitrogen of the alanine amide and the FG repeats. The P90A enhancement is specific for the pentamer-only spheres. The enhancing effects were not obvious in the GLFG phase experiments, because there, the partitioning was already complete without additional mutations. Quantifications were as in Extended Data Fig.5. **c,d,** Partitioning of cone-shaped CLPs into the FSFG phase. The stronger cohesion of this phase results in partitioning defect of wildtype CLPs, which arrest at the surface of the phase. This defect can be compensated by the G89V mutation, which allows for stronger hydrophobic capsid-FG interactions. *P*-values were calculated using unpaired two-tailed Student’s t test. *P-values*: * < 0.05, ****<0.0001. Experiments were repeated independently with identical outcomes (*N* = 3). Scale bars, 10 μm.

**Extended Data Fig. 9.**
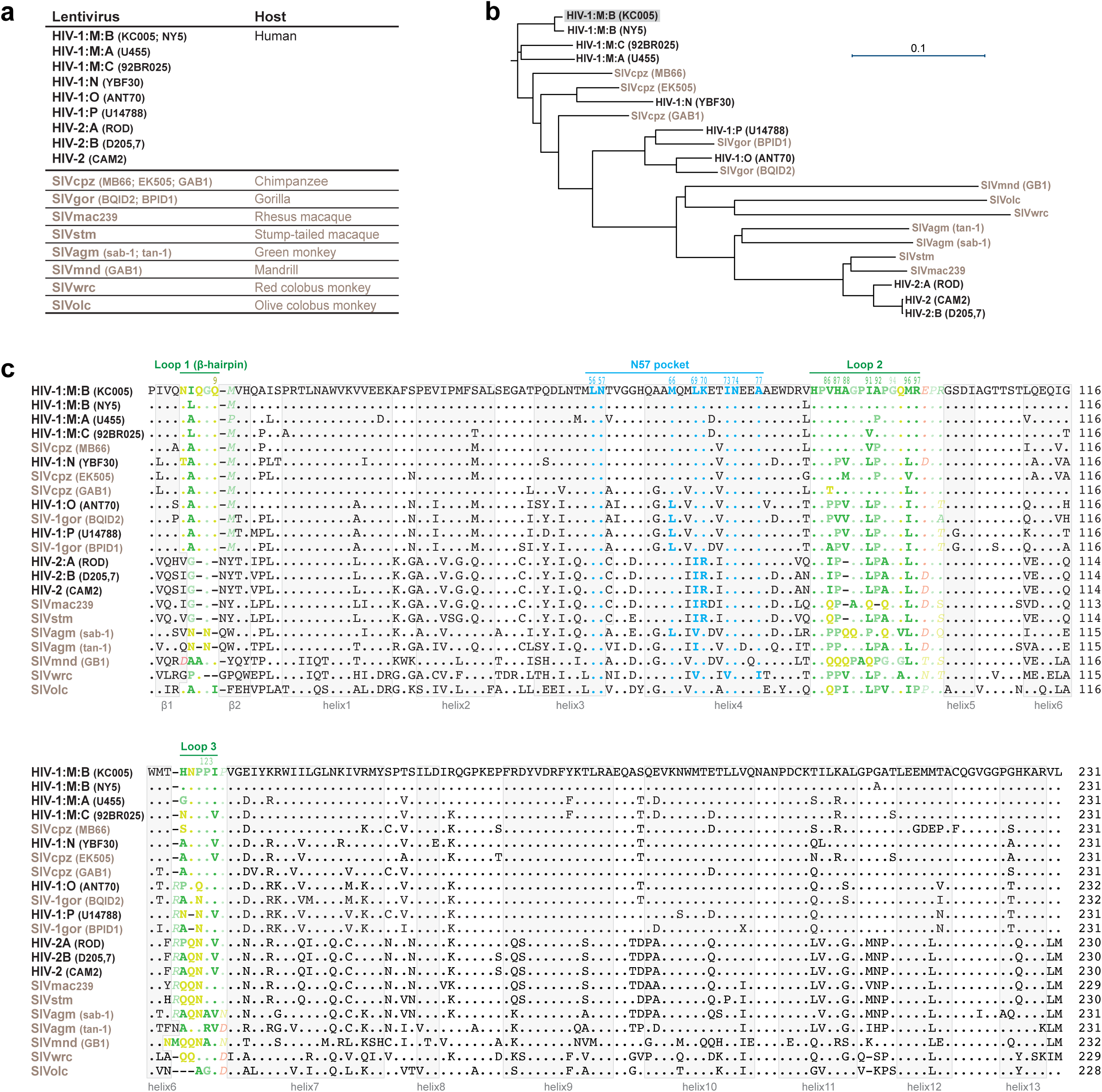
Sequence alignment and phylogenetic analysis of representative lentiviral CA proteins. **a,** Representative lentiviruses and their hosts **b,** Neighbor-joining phylogenetic tree of lentiviral CA proteins. **c,** A structural alignment of lentiviral CA sequences (based on Alphafold3-modelling). HIV-1 isolate KC005 (M group, subtype B) is the CA species analyzed in this study, therefore serving as a reference. Residues of the N57 pocket that contact the FG motif are colored in cyan. Loop1-3 residues are colored according to the amino acid scale of Fig.2a. Highly exposed residues on the outer loops are in bold. Residues that are not well exposed for FG interaction (either due to forming existing H-bond or salt bridge or due to orientation) are in italic. Secondary structures are indicated by light gray boxes. Sequence identifiers are listed in Supplementary Table S7. For sequence variations within the HIV-1: M group, see also ref.^108,109^ and Table 1.

**Extended Data Fig. 10.**
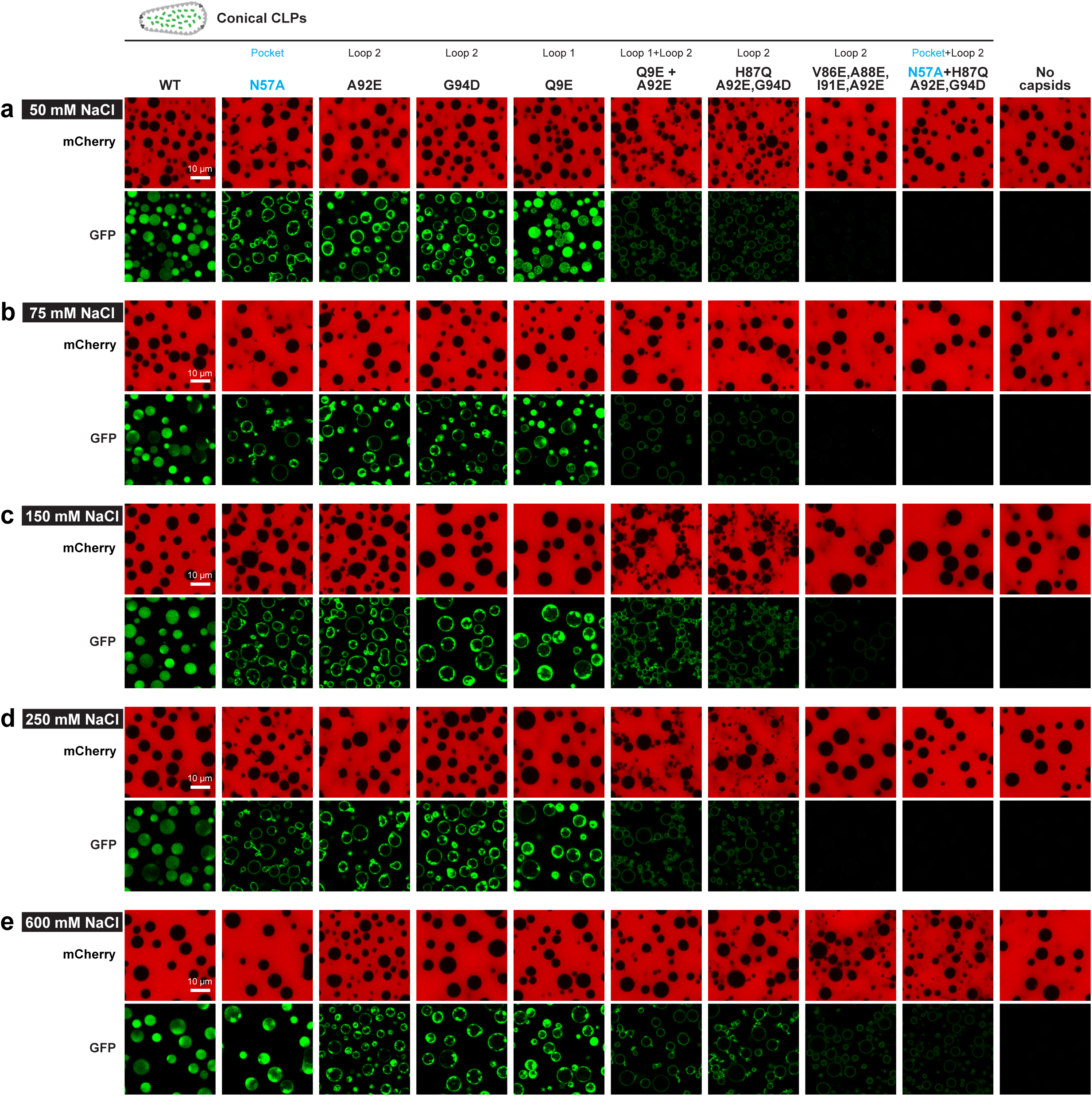
FG pocket and capsid surface residues contribute different qualities of FG interactions. Partitioning experiments into the GLFG 12mer FG phase of CLP mutations were performed as in Fig.3, but the NaCl concentration was varied from 50 mM to 600 mM as indicated in each panel. Note that wildtype CLPs showed a perfect FG phase partitioning at all salt concentrations. N57A mutant CLPs showed a strong partitioning defect at 50 mM, 75 mM, 150 mM, and 250 mM NaCl; however, at 600 mM NaCl, the N57A mutant CLPs efficiently entered the FG phase. The A92E and G94D mutations showed a strong partitioning defect at any salt concentration. The Q9E mutant CLPs entered the phase at low salt, but showed a partitioning defect at higher salt. These different behaviors indicate that N57 and the critical surface residues mediate interactions of different qualities.

## Acknowledgements

We wish to thank Thomas U. Schwartz and Onno Ackermans for fruitful collaborations, discussions, and support, Sheung Chun Ng and Jürgen Schünemann for supply with FG domains, Kathrin Gregor for the labelled xhNup133-Nb2t nanobody, Waltraud Taxer for producing the CPSF5·CPSF6 complex, Stefanie Reiter and Svetlana Agafonova for cell culture support, Susanne Brandfass and Gaby Hawlitschek for technical help, and Thomas U. Schwartz, Sheung Chun Ng, and Loren Andreas for critical reading of the manuscript.

## Supplementary information including

**Supplementary Figure S1.**
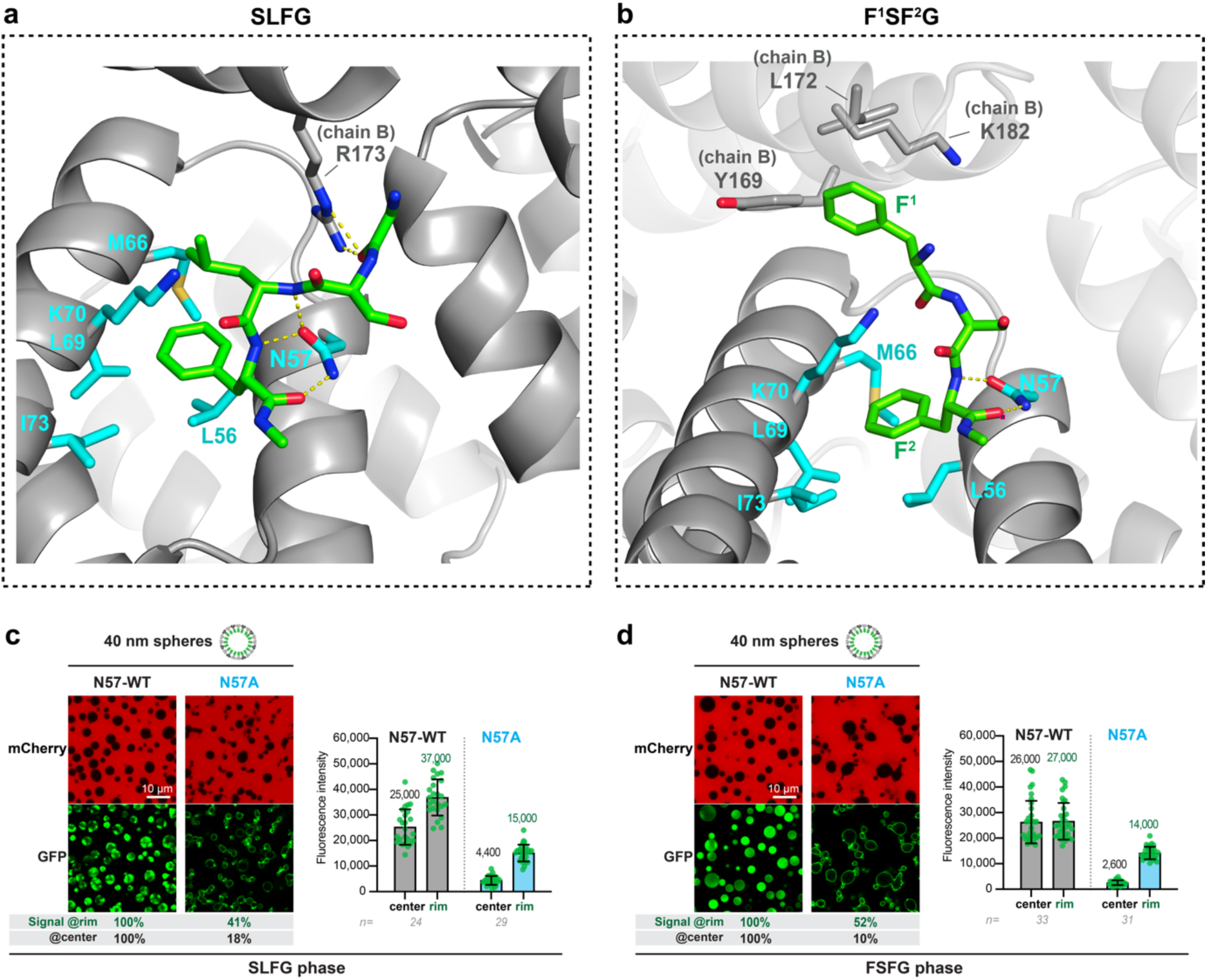
Alphafold 3-modeling of SLFG and FSFG peptides bound by the N57 pocket. **a**, Modelling of an SLFG peptide docked into the N57 pocket of an hexameric capsomer was as described in Extended Data Fig.3, the only difference being the sequence of peptide (QPATG**SLFG**GNTQ). N57 engages here in three H-bonds with the peptide backbone. For clarity, only the SLFG motif is shown. **b,** Model of a docked FSFG peptide of the sequence NTQPATG**FSFG**GNTQPATG, but only the FSFG peptide is shown. The first phenylalanine (F^1^) of the FSFG peptide makes additional hydrophobic contacts with Y169, L172, and (the aliphatic part of) K182 of chain B. **c, d,** Wildtype and N57A mutant 40 nm spheres were allowed to partition into SLFG and FSFG phases essentially as described in Extended Data Fig.3e. The N57A mutation reduced capsid-partitioning into the SLFG and the FSFG phase, validating the relevance of N57 H-bonds for general FG peptide-binding.

**Supplementary Figure S2.**
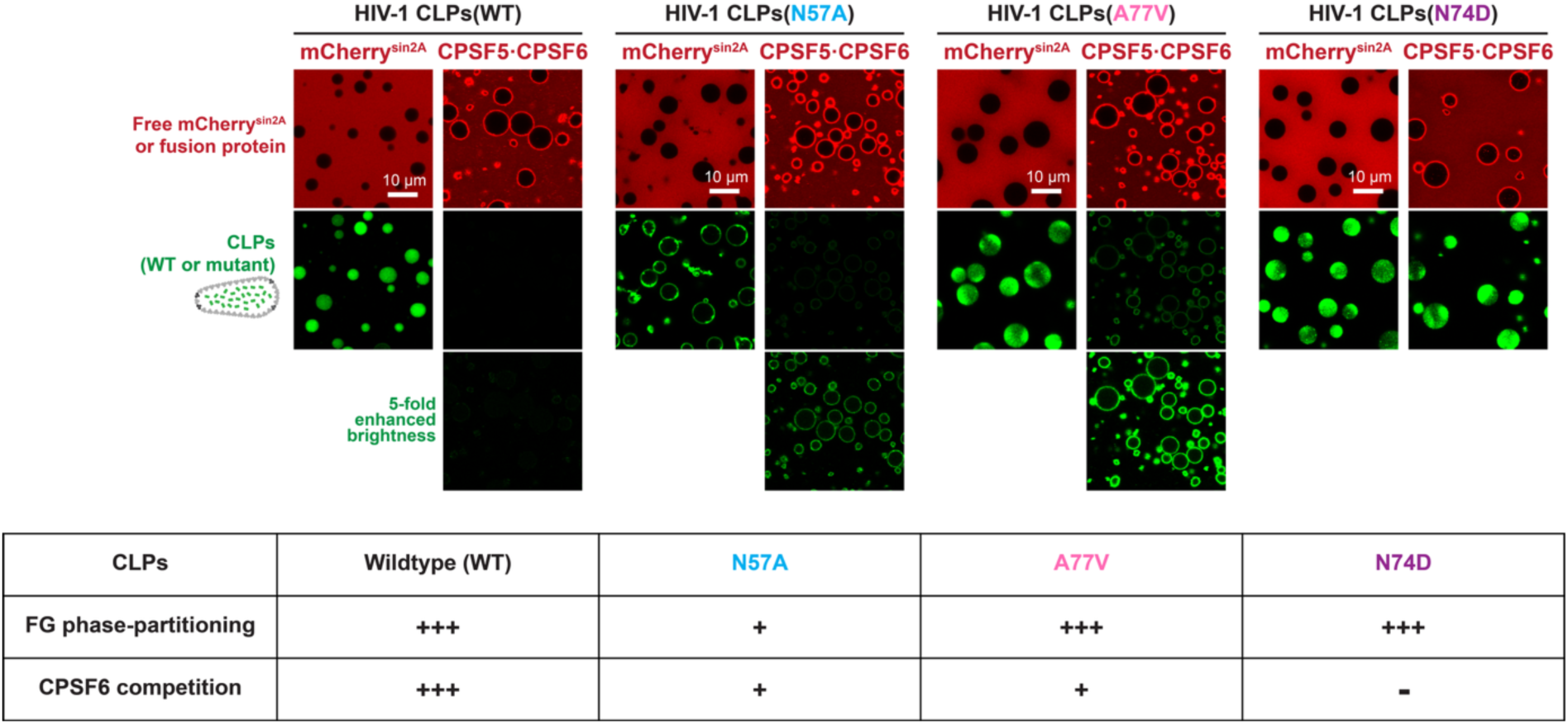
Pocket mutations discriminate between binding of GLFG repeats and the CPSF6 FG repeat. Experiment compares wildtype CLPs with CLPs that carry ‘pocket mutations’ known to interfere with the CPSF6 interaction^1-3^. Capsids were allowed to partition in an GLFG phase, either directly or in the presence of the CPSF5·CPSF6 complex (at a 2.5:1 molar ratio to CA), as described in Fig.7a. Wildtype CLPs efficiently partitioned in the GLFG phase and this partitioning was fully blocked by CPSF6 (∼0.2% residual signal). The N57A mutation impedes (but not fully abrogates) the interactions with CPSF6 and with barrier-forming FG repeats. The pentamer-only mutations (see Fig.7b) and the N74D exchange are most detrimental to the CPSF6 interaction, while causing no defect in FG phase-partitioning. A77V weakens interactions with CPSF6 but not with standard barrier-forming FG repeats. The lower tolerance to mutations indicates that the CPSF6 FG repeat docks into the pocket with more constraints and more contacts than a GLFG repeat unit of the FG phase. Also note that the CPSF5-CPSF6 complex bound prominently to the outside of the FG phase. An explanation is that the proline-rich low complexity region (PR-LCR) of CPSF6 inserts into the FG phase but the FG-repulsive CPSF6 N-terminus prevents a full entry (see Supplementary Fig.3). The capsid does not co-localize with this surface-bound CPSF6 population, probably because the PR-LCR (containing the CPSF6 FG motif) is not available for capsid-binding.

**Supplementary Figure S3.**
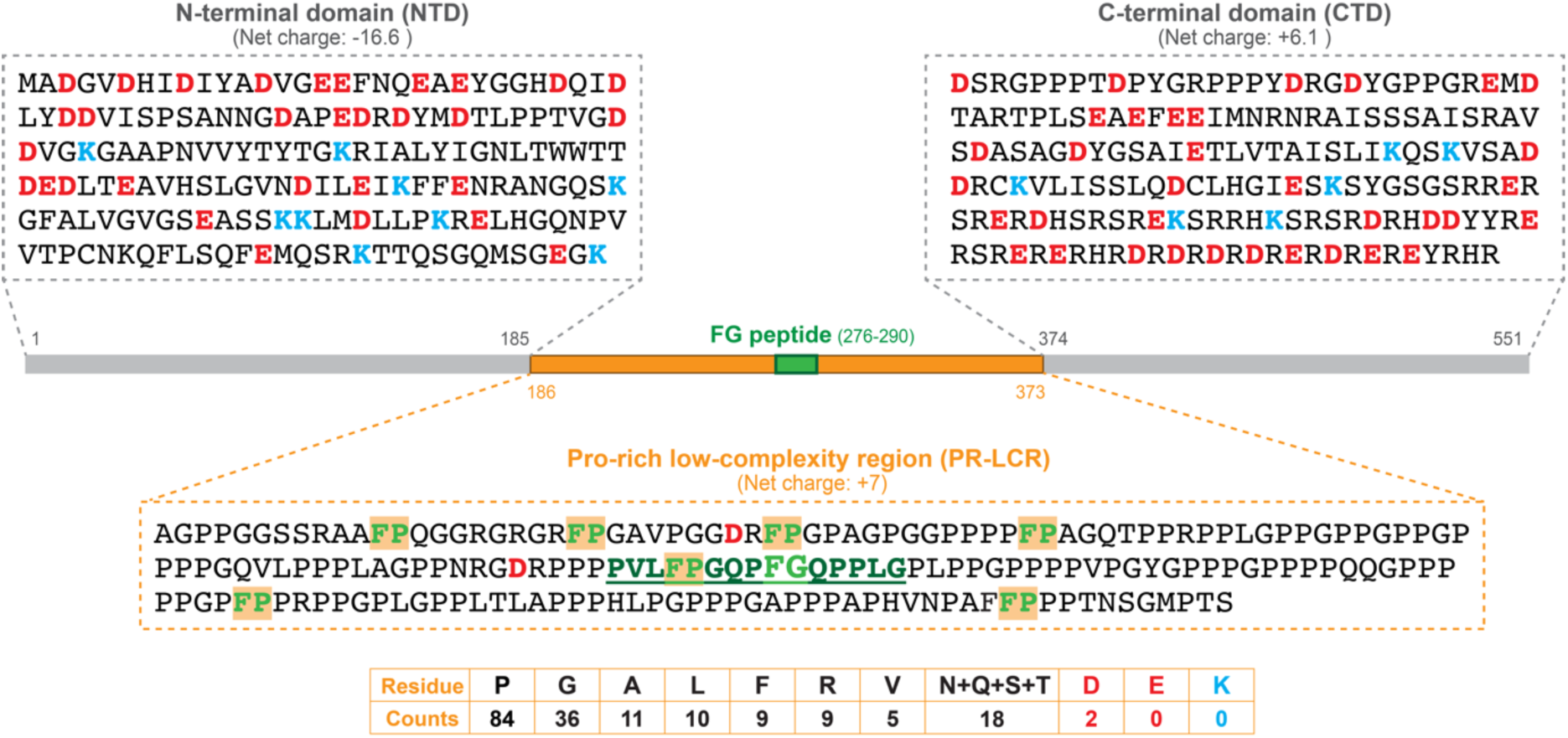
Sequence features of CPSF6 and their relation to capsid release from the FG phase. Sequence of CPSF6 (*UniProtKB:Q16630-1*), comprising three regions of different amino acid compositions:

- An N-terminal domain (NTD). It includes an RRM domain for heterotetramerization with CPSF5^4^. Its strong negative charge should make it FG-repulsive.
- A proline-rich low complexity region (PR-LCR). It lacks FG-repulsive, charged residues (K, D, E) but is rather hydrophobic, whereby not only typical hydrophobic residues (like F, L, V and A) contribute but also proline, which accounts for 45% of the residues and provides (like valine) four aliphatic carbons for hydrophobic interactions. In terms of charge-depletion and hydrophobicity, the PR-LCR resembles an Nup98 FG domain. Since 7 of the 8 phenylalanines are in an FP and not in an FG context, the PR-LCR is, however, unlikely to recruit cellular NTRs. Nevertheless, it should readily phase-separate and engage in fuzzy hydrophobic interactions with the non-charged surface of the HIV capsid, once the embedded FG motif has docked into the N57 pocket of the capsid.
- A C-terminal domain that comprises the nuclear import signal of CPSF6. It probably also can phase-separate e.g. through R-Y cation-π interactions, and this propensity gets likely increased when phosphorylation of its SR motifs shifts its net charge to zero (isoelectric precipitation). Higher levels of phosphorylation would impede condensation and render the domain FG-repulsive. This architecture can explain why CPSF6 is so effective in antagonizing the partitioning of the HIV capsid into the FG phase and releasing it from the NPC into the nucleus: The PR-LCR forms a condensate layer around the capsid, thereby masking the N57 FG-binding pocket as well as the FG-attractive surface, while the negatively charged NTD remains excluded from the local condensate and forms an outer layer with FG-repulsive properties. This model attributes a second function to the very biased surface amino acid composition of the capsid, namely not only to allow a partitioning into the FG phase but also to recruit CPSF6 as a release factor. Note that while some aspects of this model still need some testing, others are already well supported (see e.g. ref.^5-9^ and this study).

**Supplementary Table S1.**
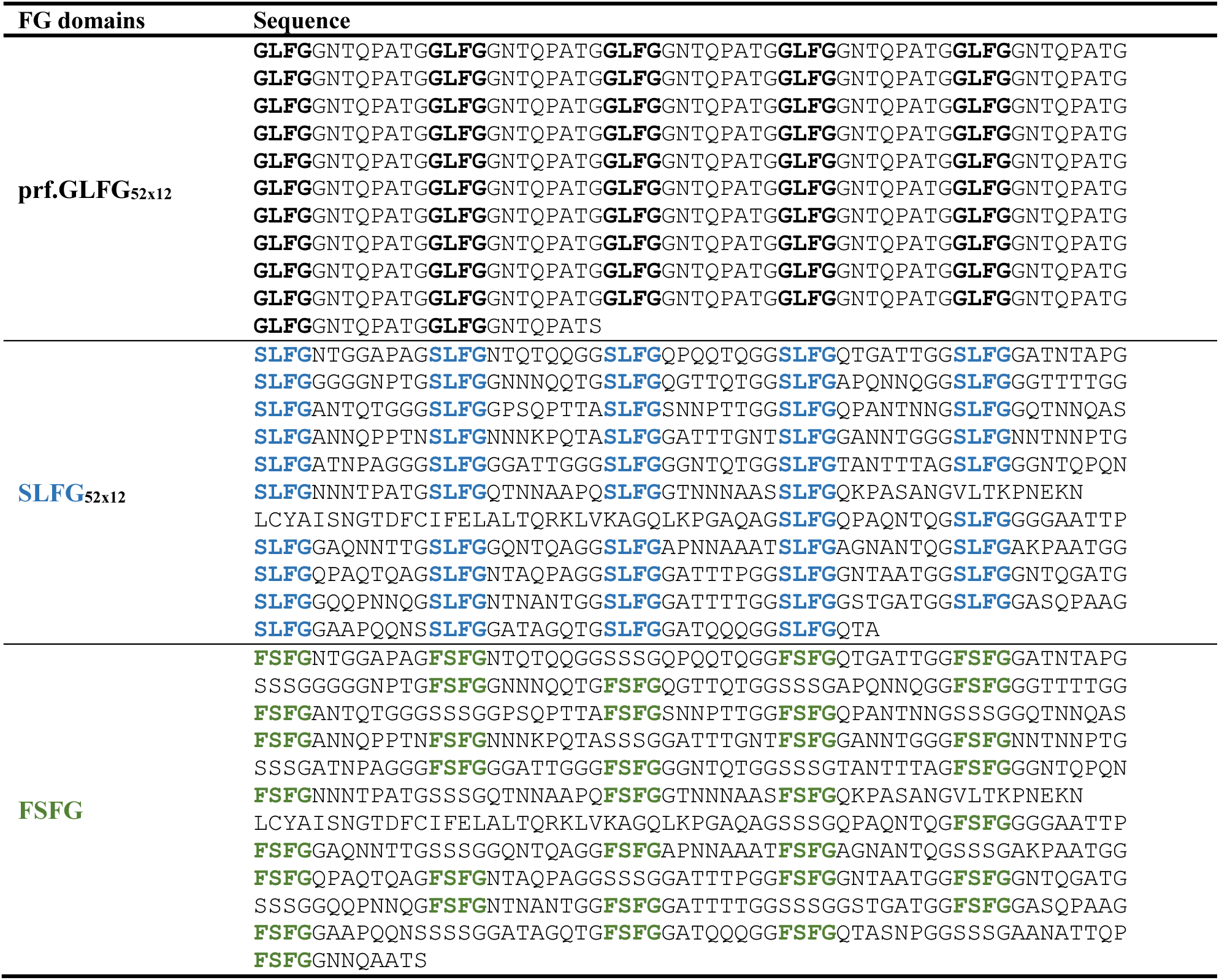
Sequences of FG domains used for FG phase experiments.

**Supplementary Table S2.** Vectors to produce HIV-1 CA proteins for assembling CLPs.

| Plasmid | CA mutation(s) | C-terminal tag | Figure | Reference |
| --- | --- | --- | --- | --- |
| pLF706 | WT | - | 1,3,4,5,6,7; ED4,5,6,7,8,10 | <sup>10</sup> |
| pLF707 | WT | EGFP | 1,6,7 | <sup>10</sup> |
| pLF974 | WT | mScarlet-I3 | 6 | This study |
| pLF721 | N57A | - | 1,3,4,5,6; ED4,5,7,10 | This study |
| pLF731 | N57A | EGFP | 1,6 | This study |
| pLF722 | N74D | - | 3; ED5 | This study |
| pLF723 | A77V | - | 3; ED5 | This study |
| pLF972 | K182R | - | 3; ED5 | This study |
| pLF856 | V86E | - | 3,5; ED4,5 | This study |
| pLF855 | V86G | - | 3; ED5 | This study |
| pLF724 | H87Q | - | 3,5; ED4,5 | This study |
| pLF859 | H87E | - | 3,5; ED4,5 | This study |
| pLF858 | H87G | - | 3; ED5 | This study |
| pLF862 | A88E | - | 3,5; ED4,5 | This study |
| pLF861 | A88G | - | 3; ED5 | This study |
| pLF725 | G89V | - | 3; ED5,8 | This study |
| pLF726 | P90A | - | 3; ED5 | This study |
| pLF867 | I91E | - | 3,5; ED4,5 | This study |
| pLF866 | I91G | - | 3; ED5 | This study |
| pLF727 | A92E | - | 3,4,5; ED4,5,6,7,10 | This study |
| pLF869 | A92G | - | 3; ED5 | This study |
| pLF728 | G94D | - | 3,4,5; ED4,5,7,10 | This study |
| pLF873 | M96E | - | 3,5; ED4,5 | This study |
| pLF872 | M96G | - | 3; ED5 | This study |
| pLF875 | R97K | - | 3,5; ED5,6 | This study |
| pLF729 | E45A | - | 3; ED5 | This study |
| pLF838 | G116A | - | 3; ED5 | This study |
| pLF966 | R132K | - | 3; ED5,6 | This study |
| pLF967 | L136M | - | 3; ED5 | This study |
| pLF969 | L136E | - | ED6 | This study |
| pLF970 | R143A | - | 3; ED5 | This study |
| pLF971 | R143K | - | ED6 | This study |
| pLF958 | Q9E | - | 3,4,5; ED4,5,7,10 | This study |
| pLF964 | P123E | - | 3,5; ED4,5 | This study |
| pLF846 | H87Q,A92E,G94D | - | 4,5; ED4,7,10 | This study |
| pLF842 | N57A+A92E | - | 4; ED4,7 | This study |
| pLF844 | N57A+G94D | - | 4; ED4,7 | This study |
| pLF956 | N57A+A92E,G94D | - | 4; ED4,7 | This study |
| pLF848 | N57A+H87Q,A92E,G94D | - | 4,5; ED4,7,10 | This study |
| pLF1018 | Q9E+A92E | - | 4,5; ED4,7,10 | This study |
| pLF930 | V86E,A88E,I91E,A92E | - | 4,5,6; ED4,7,10 | This study |
| pLF1010 | V86E,A88E,I91E,A92E | EGFP | 6 | This study |

All plasmids encode N-terminal His<sub>14</sub>-bdSUMO tagged proteins for expression in *E. coli*, and all CA proteins carry a P1A mutation to allow for tag cleavage by bdSEN1.

**Supplementary Table S3.**
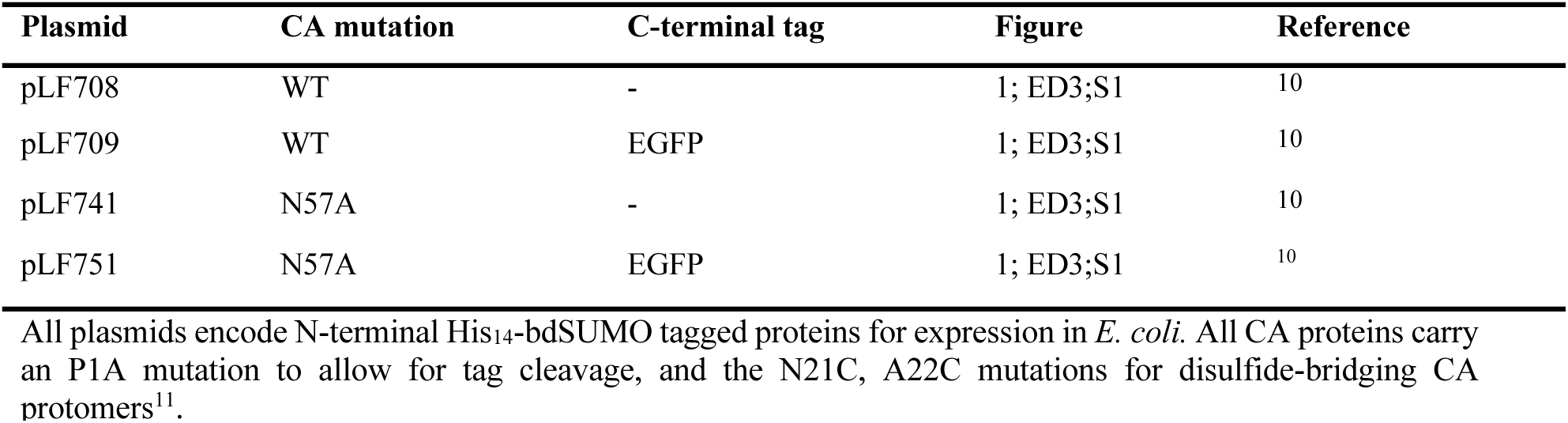
Vectors to produce CA variants for assembling 40 nm spheres.

| Plasmid | CA mutation | C-terminal tag | Figure | Reference |
| --- | --- | --- | --- | --- |
| pLF708 | WT | - | 1; ED3;S1 | 10 |
| pLF709 | WT | EGFP | 1; ED3;S1 | 10 |
| pLF741 | N57A | - | 1; ED3;S1 | 10 |
| pLF751 | N57A | EGFP | 1; ED3;S1 | 10 |

All plasmids encode N-terminal His<sub>14</sub>-bdSUMO tagged proteins for expression in *E. coli*. All CA proteins carry an P1A mutation to allow for tag cleavage, and the N21C, A22C mutations for disulfide-bridging CA protomers<sup>11</sup>.

**Supplementary Table S4.** Vectors to produce CA variants for pentamer-only 20 nm spheres.

| Plasmid | CA mutation | C-terminal tag | Figure | Reference |
| --- | --- | --- | --- | --- |
| pLF710 | WT | - | 1,4,7; ED1,4,7,8 | This study |
| pLF711 | WT | EGFP | 1,4,7; ED1,4,7,8 | This study |
| pLF761 | N57A | - | 1,4; ED4,7,8 | This study |
| pLF771 | N57A | EGFP | 1,4; ED4,7,8 | This study |
| pLF764 | H87Q | - | 4; ED4,7,8 | This study |
| pLF774 | H87Q | EGFP | 4; ED4,7,8 | This study |
| pLF765 | G89V | - | ED8 | This study |
| pLF775 | G89V | EGFP | ED8 | This study |
| pLF766 | P90A | - | ED8 | This study |
| pLF776 | P90A | EGFP | ED8 | This study |
| pLF767 | A92E | - | 4; ED4,7,8 | This study |
| pLF777 | A92E | EGFP | 4; ED4,7,8 | This study |
| pLF768 | G94D | - | 4; ED4,7,8 | This study |
| pLF778 | G94D | EGFP | 4; ED4,7,8 | This study |
| pLF990 | Q9E | - | 4; ED4,7,8 | This study |
| pLF991 | Q9E | EGFP | 4; ED4,7,8 | This study |
| pLF992 | Q9E+A92E | - | 4; ED4,7 | This study |
| pLF993 | Q9E+A92E | EGFP | 4; ED4,7 | This study |
| pLF1064 | H87Q,A92E,G94D | - | 4; ED4,7 | This study |
| pLF1065 | H87Q,A92E,G94D | EGFP | 4; ED4,7 | This study |
| pLF1066 | V86E,A88E,I91E,A92E | - | 4; ED4,7 | This study |
| pLF1067 | V86E,A88E,I91E,A92E | EGFP | 4; ED4,7 | This study |

All plasmids encode N-terminal His<sub>14</sub>-bdSUMO tagged proteins for expression in *E. coli*. All CA proteins carry an P1A mutation to allow for tag cleavage, and the G60A,G61P mutations that convert the capsid into the pentamer-only structure<sup>12</sup>.

**Supplementary Table S5.** Plasmids used for other recombinant protein expression.

| Protein | Plasmid | Encoding | Figure(s) | Reference |
| --- | --- | --- | --- | --- |
| **Probes** |  |  |  |  |
| mCherry | pDG2442 | His<sub>14</sub>-bdSUMO-mCherry | 1,3,4;<br>ED1,3,6,8,10 | 13 |
| EGFP | pSF1526 | His<sub>14</sub>-MBP-bdSUMO-EGFP | 1; ED1 | 13 |
| sinGFP4a | pDG2754 | His<sub>14</sub>-bdSUMO-sinGFP4a | 3 | 13 |
| 2xMBP-Cys | pSNG133 | His<sub>14</sub>-bdSUMO-2xMBP-Cys | 6 | This study |
| Anti-Nup133 nanobody | pDG4439 | His<sub>14</sub>-NEDD8-xhNup133-Nb2t (+2Cys) | 1,5 | 14 |
| mCherry<sup>sin2A</sup> | pDG2763 | His<sub>14</sub>-bdSUMO-mCherry<sup>sin2A</sup> | 7 | This study |
| CPSF6-mCherry<sup>sin2A</sup> | pDG5515 | 2xZpA963-bdSUMO-CPSF6-mCherry<sup>sin2A</sup>-His<sub>8</sub> | 7 | This study |
| CPSF6<sub>(FG>GG)</sub>-mCherry<sup>sin2A</sup> | pLF1275 | 2xZpA963-bdSUMO-CPSF6<sub>(FG>GG)</sub>-mCherry<sup>sin2A</sup>-His<sub>8</sub> | 7 | This study |
| CPSF7-mCherry<sup>sin2A</sup> | pDG5521 | 2xZpA963-bdSUMO-CPSF7-mCherry<sup>sin2A</sup>-His<sub>8</sub> | 7 | This study |
| CPSF5 | pDG5263 | 2xZpA963-NEDD8-CPSF5 | 7 | This study |
| **FG domains** |  |  |  |  |
| prf.GLFG<sub>52x12</sub> | pSNG57 | His<sub>18</sub>-GLFG<sub>52x12</sub> | 1,3,4,7;<br>ED3,5,6,7,10 | 15 |
| SLFG<sub>52x12</sub> | pSNG73 | His<sub>18</sub>-SLFG<sub>52x12</sub> | ED1;S1 | 16 |
| FSFG<sub>52x12</sub> | pSNG150 | His<sub>18</sub>-FSFG domain | ED1,8;S1 | 16 |

**Supplementary Table S6.** Amino acid sequences of Nup358 from various species.

| Species | Classification | Sequence identifier |
| --- | --- | --- |
| Human<br>(*Homo sapiens*) | Class: Mammalia, Order: Primates | UniProtKB/Swiss-Prot: P49792 |
| Mouse<br>(*Mus musculus*) | Class: Mammalia, Order: Rodentia | UniProtKB/Swiss-Prot: Q9ERU9 |
| Chicken<br>(*Gallus gallus*) | Class: Aves, Order: Galliformes | GenBank: XP_040513539.1 |
| Snake<br>(*Python bivittatus*) | Class: Reptilia, Order: Squamata | GenBank: XP_007427690.1 |
| Fish<br>(*Danio rerio*) | Class: Actinopterygii, Order: Cypriniformes, | GenBank: XP_021334470.1 |
| Frog<br>(*Xenopus laevis*) | Class: Amphibia, Order: Anura | GenBank: XP_041438290.1 |
| Frog<br>(*Xenopus tropicalis*) | Class: Amphibia, Order: Anura | GenBank: AAI57560.1 |
| Frog<br>(*Rana temporaria*) | Class: Amphibia, Order: Anura | GenBank: XP_040192403.1 |
| Toad<br>(*Bufo bufo*) | Class: Amphibia, Order: Anura | GenBank: XP_040281002.1 |
| Toad<br>(*Bufo gargarizans*) | Class: Amphibia, Order: Anura | GenBank: XP_044139501.1 |
| Toad<br>(*Bombina bombina*) | Class: Amphibia, Order: Anura | GenBank: XP_053561416.1 |
| Salamander<br>(*Ambystoma mexicanum*) | Class: Amphibia, Order: Urodela | GenBank: XP_069492978.1 |
| Salamander<br>(*Pleurodeles waltl*) | Class: Amphibia, Order: Urodela | GenBank: XP_069060581.1 |
| Caecilian<br>(*Geotrypetes seraphini*) | Class: Amphibia, Order: Gymnophiona | GenBank: XP_033804751.1 |
| Caecilian<br>(*Rhinatrema bivittatum*) | Class: Amphibia, Order: Gymnophiona | GenBank: XP_029460448.1 |

**Supplementary Table S7.** Amino acid sequences of CA proteins from various lentiviruses.

| Lentivirus | Subtype | Isolate | Host | Sequence identifier |
| --- | --- | --- | --- | --- |
| **HIV-1** |  |  |  |  |
| HIV-1:M:B (KC005) | M, B | KC005 | Human<br>(*Homo sapiens*) | GenBank: CBI62787.1 |
| HIV-1:M:B (NY5) | M, B | NY5 | Human<br>(*Homo sapiens*) | GenBank: AAB04036.1 |
| HIV-1:M:A (U455) | M, A | U455 | Human<br>(*Homo sapiens*) | GenBank: AAA75018.1 |
| HIV-1:M:C (92BR025) | M, C | 92BR025 | Human<br>(*Homo sapiens*) | UniProtKB/Swiss-Prot: O12158.2 |
| HIV-1:N (YBF30) | N | YBF30 | Human<br>(*Homo sapiens*) | UniProtKB/Swiss-Prot: O91079.3 |
| HIV-1:O (ANT70) | O | ANT70 | Human<br>(*Homo sapiens*) | UniProtKB/Swiss-Prot: Q77372.3 |
| HIV-1:P (U14788) | P | U14788 | Human<br>(*Homo sapiens*) | GenBank: ADR03143.1 |
| **HIV-2** |  |  |  |  |
| HIV-2:A (ROD) | A | ROD | Human<br>(*Homo sapiens*) | UniProtKB/Swiss-Prot: P04590.3 |
| HIV-2:B (D205,7) | B | D205,7 | Human<br>(*Homo sapiens*) | UniProtKB/Swiss-Prot: P15832.3 |
| HIV-2 (CAM2) | other | CAM2 | Human<br>(*Homo sapiens*) | UniProtKB/Swiss-Prot: P24106.3 |
| **SIV** |  |  |  |  |
| SIVcpz (MB66) |  | MB66 | Chimpanzee<br>(*Pan troglodytes troglodytes*) | UniProtKB/Swiss-Prot: Q1A268.3 |
| SIVcpz (EK505) |  | EK505 | Chimpanzee<br>(*Pan troglodytes troglodytes*) | UniProtKB/Swiss-Prot: Q1A249.3 |
| SIVcpz (GAB1) |  | GAB1 | Chimpanzee<br>(*Pan troglodytes troglodytes*) | UniProtKB/Swiss-Prot: P17282.1 |
| SIVgor (BPID1) |  | BPID1 | Gorilla<br>(*Gorilla gorilla gorilla*) | GenBank: AJT49601.1 |
| SIVgor (BQID2) |  | BQID2 | Gorilla<br>(*Gorilla gorilla gorilla*) | GenBank: AJT49619.1 |
| SIVmac239 |  | SIVmac239 | Rhesus macaque<br>(*Macaca mulatta*) | GenBank: AQY10339.1 |
| SIVstm |  | STM | Stump-tailed macaque<br>(*Macaca arctoides*) | UniProtKB/Swiss-Prot: P31634.1 |
| SIVagm (sab-1) |  | sab-1 | Green monkey<br>(*Chlorocebus sabaeus*) | GenBank: AAA21504.1 |
| SIVagm (tan-1) |  | tan-1 | Tantalus monkey<br>(*Chlorocebus tantalus*) | GenBank: AAC57051.1 |
| SIVmnd-2 |  | SIVmnd-2 | Mandrill<br>(*Mandrillus sphinx*) | GenBank: AAM08705.2 |
| SIVmnd (GB1) |  | GB1 | Mandrill<br>(*Mandrillus sphinx*) | UniProtKB/Swiss-Prot: P22381.1 |
| SIVwrc |  | WRC | Red colobus monkey<br>(*Piliocolobus badius temminckii*) | GenBank: CAP72503.1 |
| SIVolc |  | OLC | Olive colobus monkey<br>(*Procolobus verus*) | GenBank: CAQ60124.1 |

